# Lamin A/C coordinates nuclear mechanics, chromatin architecture, and transcriptional homeostasis in skeletal muscle *in vivo*

**DOI:** 10.64898/2026.05.31.729137

**Authors:** Soham Ghosh, Kanita Hrustanovic, Stephanie E. Schneider, Adrienne K. Scott, Jessica Kelly, Natalie Calahan, Benjamin Seelbinder, Brittany M. St Martin, Xin Xu, Corey P. Neu

## Abstract

The nuclear lamina provides mechanical integrity to the eukaryotic nucleus and organizes lamina-associated chromatin domains that are important for chromatin architecture and gene-expression regulation. Lamin A/C, a major component of the nuclear lamina, is disrupted in hereditary laminopathies and has also been implicated in aging-associated nuclear dysfunction. Although the mechanical role of Lamin A/C has been extensively studied *in vitro*, particularly in monolayer cell culture and isolated nuclei, its role in maintaining nuclear mechanics and chromatin organization in intact tissues remains incompletely understood. Here, we investigated how partial and complete Lamin A/C disruption affects nuclear shape, chromatin architecture, intranuclear mechanics, and gene expression *in vivo*. Across multiple murine tissues, Lamin A/C deficiency did not cause a generalized collapse of nuclear shape or gross tissue architecture. Instead, Lamin A/C disruption preferentially altered chromatin architecture in mechanically stiff tissues, including skeletal muscle and heart. Using live *in vivo* deformation microscopy during controlled hindlimb muscle stimulation, we quantified real-time multiscale deformation of skeletal muscle tissue and nuclei. These measurements revealed reduced effective nuclear stiffness and altered load sharing between euchromatin-rich and heterochromatin-rich domains after Lamin A/C loss. Super-resolution imaging further showed that partial and complete Lamin A/C disruption uncoupled H3K9me3 from DAPI-dense heterochromatin, indicating a spatial disruption of repressive chromatin organization. Exploratory ATAC-seq suggested increased chromatin accessibility in heterozygous muscle, whereas RNA-seq showed that complete Lamin A/C loss caused broad myopathic transcriptional dysregulation while partial loss preserved a near-wild-type transcriptomic state. Integrated analysis identified HDAC2 as a candidate mechanosensitive compensatory node that may help buffer gene expression after partial Lamin A/C disruption. Together, these results establish Lamin A/C as an *in vivo* coordinator of nuclear mechanics, heterochromatin organization, and transcriptional homeostasis in skeletal muscle.

## INTRODUCTION

Tissue phenotype is defined by composition, architecture, and biomechanics. These properties depend on the expression and organization of proteins in the cell, and ultimately on biochemical activity in the nucleus. Conversely, the mechanical properties and forces generated by tissues are transmitted back to the nucleus through the extracellular matrix, cytoskeleton, and nuclear envelope, where they influence transcription factors, chromatin organization, and epigenetic mechanisms. This reciprocal communication between tissue phenotype and gene expression is a central feature of multiscale mechanobiology and depends on mechanotransducers that connect physical forces to nuclear regulation^1^. Several structures at the cell membrane, cytoskeleton, and nucleus have emerged as mechanosensitive regulators of gene expression. Among these, the LINC complex and the nuclear lamina are of special interest because they provide a physical and biochemical connection between the cytoskeleton and chromatin^2^.

The nuclear lamina has been studied extensively for more than two decades as both a structural scaffold and a regulator of nuclear function^3,4^. High-resolution structural studies have shown that the lamina is a mesh-like network composed primarily of A-type lamins, Lamin A and Lamin C encoded by the *LMNA* gene (*Lmna* in mice), and B-type lamins, Lamin B1 and Lamin B2 encoded by *LMNB1* and *LMNB2* (*Lmnb1* and *Lmnb2* in mice). A-type and B-type lamins form distinct but interacting networks, and they contribute differently to nuclear mechanics, nuclear stability, chromatin organization, and gene regulation^3,5^. *In vitro* studies have been particularly important in defining these mechanical roles. They showed that Lamin A/C is a major determinant of nuclear elasticity and stiffness, whereas B-type lamins contribute to other aspects of nuclear mechanical behavior and nuclear integrity^5–8^. Lamin A/C-deficient cells have softer nuclei, increased nuclear deformation under applied mechanical load, and defective mechanotransduction, including altered gene expression^6^. These findings created the foundation for the current view that Lamin A/C protects the nucleus from mechanical stress and helps convert physical forces into transcriptional responses.

In parallel, molecular and structural biology studies have shown that the nuclear lamina is not only a mechanical structure but also an organizer of chromatin^5^. Lamin-associated proteins connect the lamina to chromatin and help define lamina-associated domains, or LADs, which are generally enriched in transcriptionally repressed chromatin^9,10^. Through these interactions, Lamin A/C is positioned to regulate both the physical architecture of the nucleus and the epigenetic state of the genome^11^. Recent reviews have emphasized that lamins act at the intersection of nuclear mechanics, chromatin organization, and transcriptional regulation, especially in mechanically active tissues and in laminopathies^12–14^.

Despite this extensive literature, a key gap remains. Most mechanistic studies of Lamin A/C have been performed in cultured cells, isolated nuclei, or simplified *in vitro* systems. These approaches are powerful because they allow direct manipulation and measurement of nuclear mechanics, but they do not fully reproduce the three-dimensional and mechanically constrained environment of intact tissues. This limitation is especially important in mechanobiology. Cell shape, substrate stiffness, extracellular matrix structure, cytoskeletal tension, and tissue-level force transmission all influence nuclear morphology and chromatin organization. Cells cultured as two-dimensional monolayers can therefore reveal important cell-autonomous phenotypes, but they may also exaggerate or redirect mechanical behavior that would be buffered differently *in vivo*^15,16^. Therefore, it remains unclear whether the classical *in vitro* phenotypes of Lamin A/C deficiency, including abnormal nuclear shape, increased nuclear deformation, and altered chromatin organization, are reproduced in intact tissues.

This question is particularly important because Lamin A/C mutations cause laminopathies^17–20^, a diverse group of diseases that often affect mechanically active tissues such as skeletal muscle, heart, bone, and adipose tissue^21,22^. The tissue specificity of laminopathies has been difficult to explain because Lamin A/C is broadly expressed and has many nuclear functions. One influential model is that tissues exposed to greater mechanical stress depend more strongly on Lamin A/C for nuclear mechanical stability. This idea is supported by the observation that the ratio of Lamin A/C to Lamin B scales with tissue stiffness^23^. However, direct *in vivo* evidence linking Lamin A/C deficiency to altered nuclear mechanics, chromatin architecture, and gene expression inside mechanically active tissues remains limited. The field has therefore relied heavily on extrapolating from *in vitro* systems to infer how Lamin A/C functions *in vivo*.

Intranuclear chromatin mechanics adds another layer of complexity. Chromatin is not only a substrate for transcription and epigenetic regulation; it also contributes to the mechanical behavior of the nucleus^24–26^. Recent work has shown that nuclear deformation can reorganize chromatin and influence cell phenotype, suggesting that chromatin architecture itself can act as a mechanosensitive component of the nucleus. Therefore, the role of Lamin A/C *in vivo* may not be limited to maintaining the nuclear outline. Lamin A/C may also coordinate how mechanical forces are distributed across chromatin domains, how repressive chromatin marks remain spatially organized, and how gene expression remains stable in mechanically loaded tissues.

This possibility is also relevant beyond rare laminopathies. Lamin A/C biology has been connected to progeroid syndromes, nuclear defects in aging cells, altered chromatin organization, DNA damage responses, and tissue degeneration^27–29^. However, aging is complex and cannot be reduced to Lamin A/C deficiency alone. A more balanced view is that Lamin A/C-dependent nuclear mechanics and chromatin organization may represent one mechanism by which mechanically active tissues maintain nuclear homeostasis over time. Understanding this mechanism *in vivo* may therefore provide insight not only into laminopathy, but also into broader questions of tissue maintenance, degeneration, and aging-associated nuclear dysfunction.

In this study, we aimed to close this knowledge gap by investigating the mechanobiological role of Lamin A/C in intact tissues and live skeletal muscle. Quantifying nuclear mechanics *in vivo* is technically challenging because the nucleus must be imaged and mechanically loaded within its native tissue environment. In previous work, we established an image-based deformation microscopy approach that allows intranuclear mechanics to be quantified in live mouse skeletal muscle with high spatial resolution during controlled muscle stimulation^30,31^. Here, we used this approach together with tissue imaging, chromatin architecture analysis, Airyscan super-resolution microscopy, ATAC-seq, and RNA-seq to investigate how Lamin A/C disruption affects nuclear shape, chromatin organization, nuclear mechanics, and gene expression *in vivo*.

We used a Lamin A/C mutant mouse model in which Lamin A/C does not properly integrate into the nuclear lamina, comparing wild-type (*Lmna*^+/+^; WT), heterozygous (*Lmna*^+/-^; HZ), and Lamin A/C-deficient (*Lmna*^-/-^; LD) mice. We first asked whether the expected nuclear-shape phenotype of Lamin A/C deficiency is present across intact tissues. We then examined tissue architecture and chromatin organization across tissues with different mechanical environments. Next, we used live *in vivo* confocal deformation microscopy to quantify tissue-to-nucleus strain transfer and intranuclear strain distribution in skeletal muscle. Finally, we integrated super-resolution imaging of chromatin marks with chromatin-accessibility and transcriptomic profiling to identify how partial and complete Lamin A/C disruption affect chromatin regulation and transcriptional homeostasis.

The results reveal that Lamin A/C deficiency does not primarily appear *in vivo* as a generalized collapse of nuclear shape. Instead, Lamin A/C disruption produces a more internal and tissue-context-dependent phenotype: altered chromatin architecture in mechanically stiff tissues, reduced effective nuclear stiffness in skeletal muscle, disrupted load sharing between euchromatin and heterochromatin, uncoupling of H3K9me3 from DAPI-dense heterochromatin, and nonlinear transcriptional responses to Lamin A/C dosage. Partial Lamin A/C disruption produces measurable mechanical and epigenetic remodeling while preserving a near-WT transcriptomic state, suggesting compensatory buffering. Complete Lamin A/C loss exceeds this buffering capacity and leads to broad myopathic transcriptional dysregulation. Together, these findings support a model in which Lamin A/C maintains mechano-epigenetic homeostasis *in vivo* by coordinating nuclear mechanics, chromatin architecture, and gene-expression stability in skeletal muscle.

## RESULTS

### Lamin A/C deficiency does not substantially alter nuclear shape in intact tissues in vivo

To investigate how Lamin A/C deficiency affects nuclear organization in its native physiological environment, we first examined nuclear shape across multiple murine tissues *in vivo* (Figure 1). Lamin A/C has long been considered a major determinant of nuclear stiffness and nuclear shape, largely based on studies in cells cultured on two-dimensional substrates. Consistent with those previous observations^6^, we found that skin-derived fibroblasts and cardiac fibroblasts cultured as 2D monolayers showed a subpopulation of Lamin A/C-deficient nuclei with irregular shape and occasional blebbing (Figure S2). In contrast, nuclei from wild-type (*Lmna*^+/+^; WT) and heterozygous (*Lmna*^+/-^; HZ) fibroblasts were mostly round and regular. Although this trend was evident visually, the difference in nuclear irregularity was not statistically significant, indicating that even in culture the abnormal nuclear-shape phenotype was present only in a subset of Lamin A/C-deficient (*Lmna*^-/-^; LD) cells.

**Figure 1.**
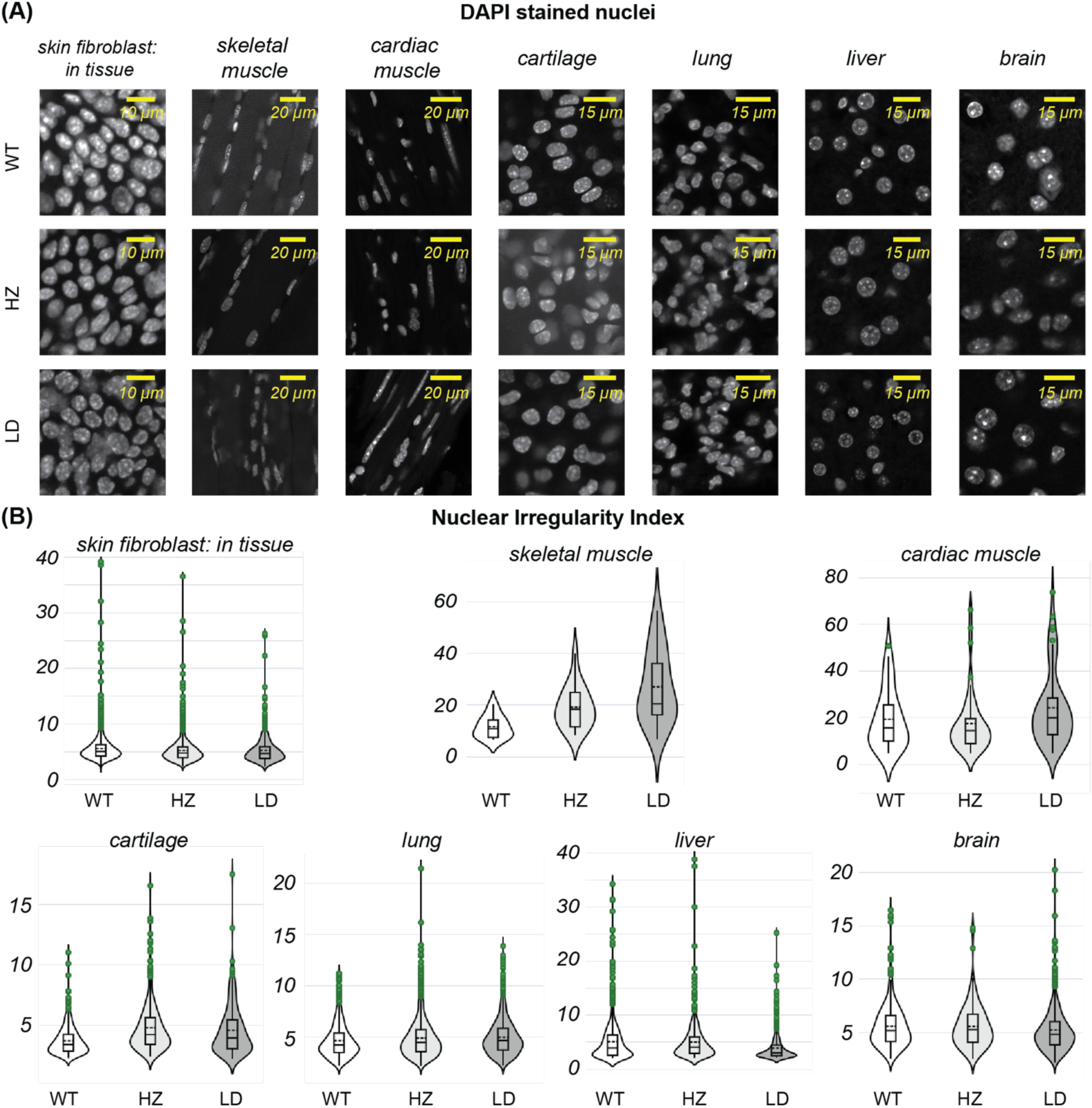
Lamin A/C deficiency does not substantially alter nuclear shape in intact tissues in vivo. **(A)** Representative confocal microscopy images showing nuclear morphology across multiple murine tissues from wild-type (WT; Lmna+/+), heterozygous (HZ; Lmna+/-), and Lamin A/C-deficient (LD; Lmna-/-) mice. Lamin A/C deficiency caused modest nuclear irregularity in skeletal muscle (tibialis anterior) and cardiac muscle nuclei, but this phenotype was not broadly observed across other tissues, including skin, cartilage, lung, liver, and brain. **(B)** Quantification of nuclear morphology using a nuclear irregularity index calculated from multiple geometric parameters, including aspect ratio, radius ratio, circularity, and nuclear area relative to bounding-box area (see Figure S1). Higher values indicate greater deviation from a regular nuclear shape. Although skeletal muscle and heart showed a modest increase in nuclear irregularity in LD mice, differences were not statistically significant. Data are based on >500 nuclei per group per tissue from at least three animals.

We next asked whether this phenotype was preserved in intact tissues, where cells and nuclei remain embedded within their native extracellular matrix and tissue architecture. Surprisingly, Lamin A/C deficiency did not result in a substantial change in nuclear shape in most tissues examined (Figure 1A). In native skin tissue, where fibroblasts showed irregular nuclear morphology in monolayer culture, nuclear shape was largely maintained *in vivo*. Similarly, connective and soft tissues including cartilage, liver, lung, and brain did not show obvious Lamin A/C-dependent changes in nuclear morphology. Skeletal muscle and cardiac muscle, which are among the most clinically affected tissues in Lamin A/C-associated disease, showed only a modest increase in nuclear irregularity in the Lamin A/C-deficient group (Figure 1B). However, this effect was not statistically significant when quantified using a nuclear irregularity index based on multiple geometric parameters (Figure S1).

These results suggest that Lamin A/C deficiency does not cause a generalized collapse of nuclear shape in intact tissues *in vivo*. Instead, the nuclear-shape phenotype appears to depend strongly on cellular context. In 2D culture, where cells spread autonomously on a rigid surface, loss of Lamin A/C can produce irregular nuclei in a subset of cells. In native tissue, however, the surrounding tissue architecture may mechanically constrain nuclear morphology and preserve gross nuclear shape, even in tissues such as skeletal muscle and heart. Thus, nuclear shape alone is not a sufficiently sensitive readout of Lamin A/C dysfunction *in vivo*, motivating us to examine whether more subtle changes occur inside the nucleus.

### Preserved tissue architecture explains the maintenance of nuclear shape in Lamin A/C-deficient tissues

Because nuclear shape is strongly influenced by cell shape, cytoskeletal organization, and the mechanical constraints imposed by the surrounding tissue^32^, we next asked whether the preserved nuclear morphology in Lamin A/C-deficient tissues was accompanied by preserved tissue-level architecture. This question is particularly important *in vivo*, where nuclei do not deform in isolation but remain mechanically integrated with the cytoskeleton, neighboring cells, and extracellular matrix.

To assess this, we visualized F-actin organization using phalloidin staining together with Lamin B staining to identify nuclei across multiple tissues (Figure 2, Figure S3, Figure S4). In skeletal muscle and cardiac muscle, which are among the most mechanically active and clinically affected tissues in Lamin A/C-associated disease, the overall tissue architecture was largely maintained in the Lamin A/C-deficient group. Sarcomeric striations remained clearly visible, and the gross organization of muscle fibers did not show an obvious disruption compared to WT and HZ tissues. Lamin B staining further confirmed that nuclear outlines were broadly preserved, consistent with the nuclear-shape analysis described above.

**Figure 2.**
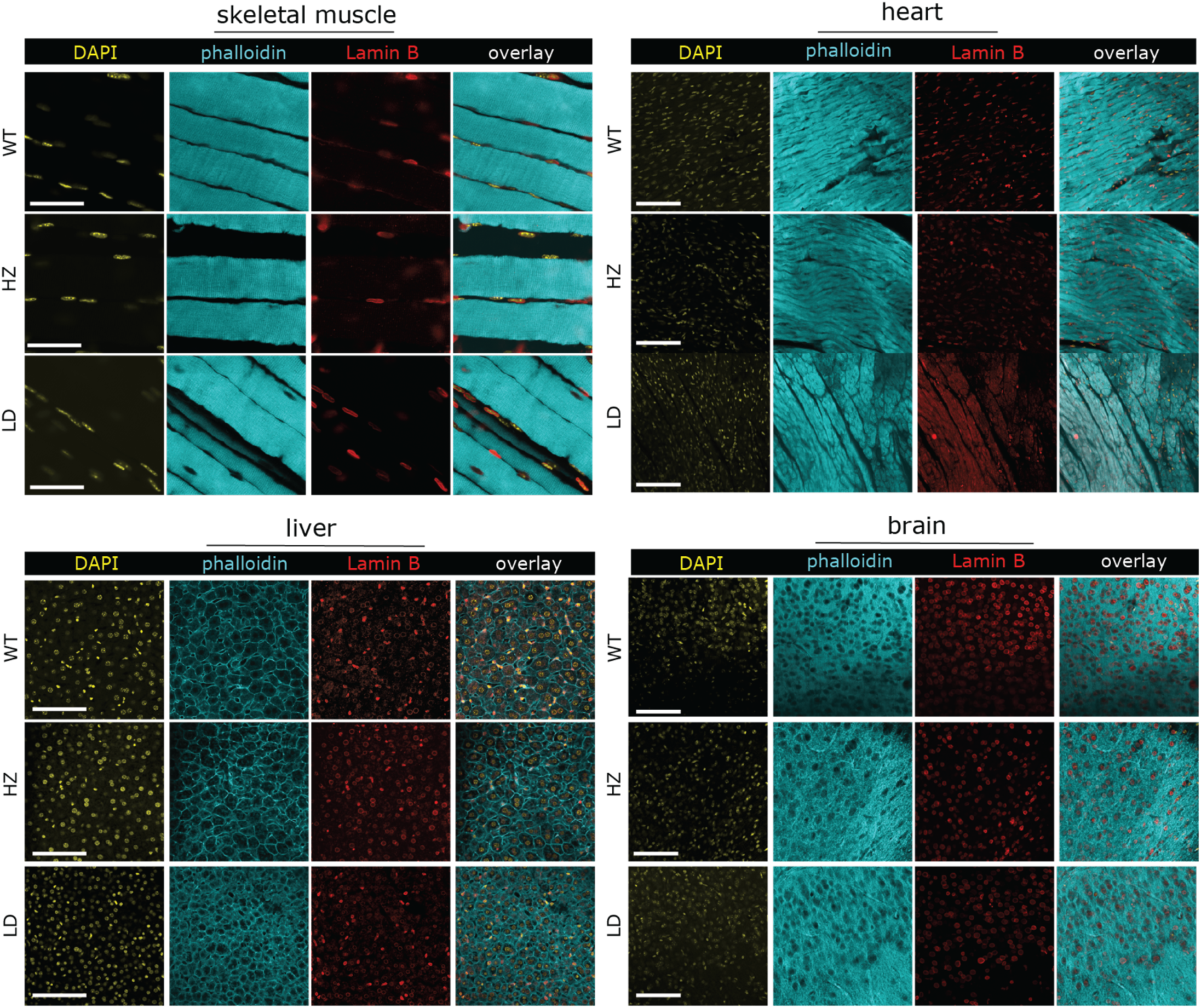
Tissue- and cytoskeletal-level architecture is grossly preserved after Lamin A/C deficiency. Phalloidin staining of F-actin was used to visualize cytoskeletal and tissue-level organization across tissues from WT, HZ, and LD mice. Confocal microscopy was used. In skeletal muscle (tibialis anterior) and heart, the tissues most clinically affected by Lamin A/C deficiency, muscle architecture remained grossly preserved, with visible striations and no obvious disruption of tissue organization. Softer tissues, including liver and brain, also showed no discernible architectural disruption. Lamin B staining further confirmed that nuclear outlines were broadly maintained across genotypes. Together, these data suggest that intact tissue architecture may help preserve gross nuclear morphology in vivo despite Lamin A/C deficiency. All scale bars: 100 µm.

A similar preservation of tissue organization was observed in other tissues. Stiffer tissues such as skin and cartilage did not show a discernible loss of tissue architecture, and softer tissues including lung, liver, and brain also appeared grossly comparable across genotypes. Although this analysis does not exclude molecular or ultrastructural defects, it indicates that Lamin A/C deficiency does not produce a broad collapse of cytoskeletal or tissue-level organization in the tissues examined.

Together with the nuclear morphology data, these findings suggest that the compressive force inside the intact tissue architecture may buffer the effect of Lamin A/C deficiency on nuclear shape *in vivo* which would otherwise demonstrate an irregular shape under predominantly mechanical tension when cultured in vitro. In native tissues, cells and nuclei are mechanically constrained within a multiscale structural environment, which may explain why the dramatic nuclear-shape abnormalities often observed in 2D culture are not reproduced as a dominant phenotype *in vivo*. This preservation of nuclear and tissue architecture led us to ask whether Lamin A/C deficiency instead alters the internal organization of the nucleus, particularly chromatin architecture.

### Chromatin architecture, rather than nuclear shape, is disrupted by Lamin A/C deficiency in mechanically stiff tissues

Because Lamin A/C deficiency did not substantially alter nuclear shape or gross tissue architecture *in vivo*, we next asked whether the internal organization of the nucleus was affected. We focused on chromatin architecture because Lamin A/C is closely associated with heterochromatin organization at the nuclear periphery, and because chromatin itself can contribute to nuclear mechanics and force transmission^25^. We therefore analyzed the spatial distribution of DAPI intensity inside individual nuclei across multiple tissues, imaged using laser scanning confocal microscope (Figure 3).

**Figure 3.**
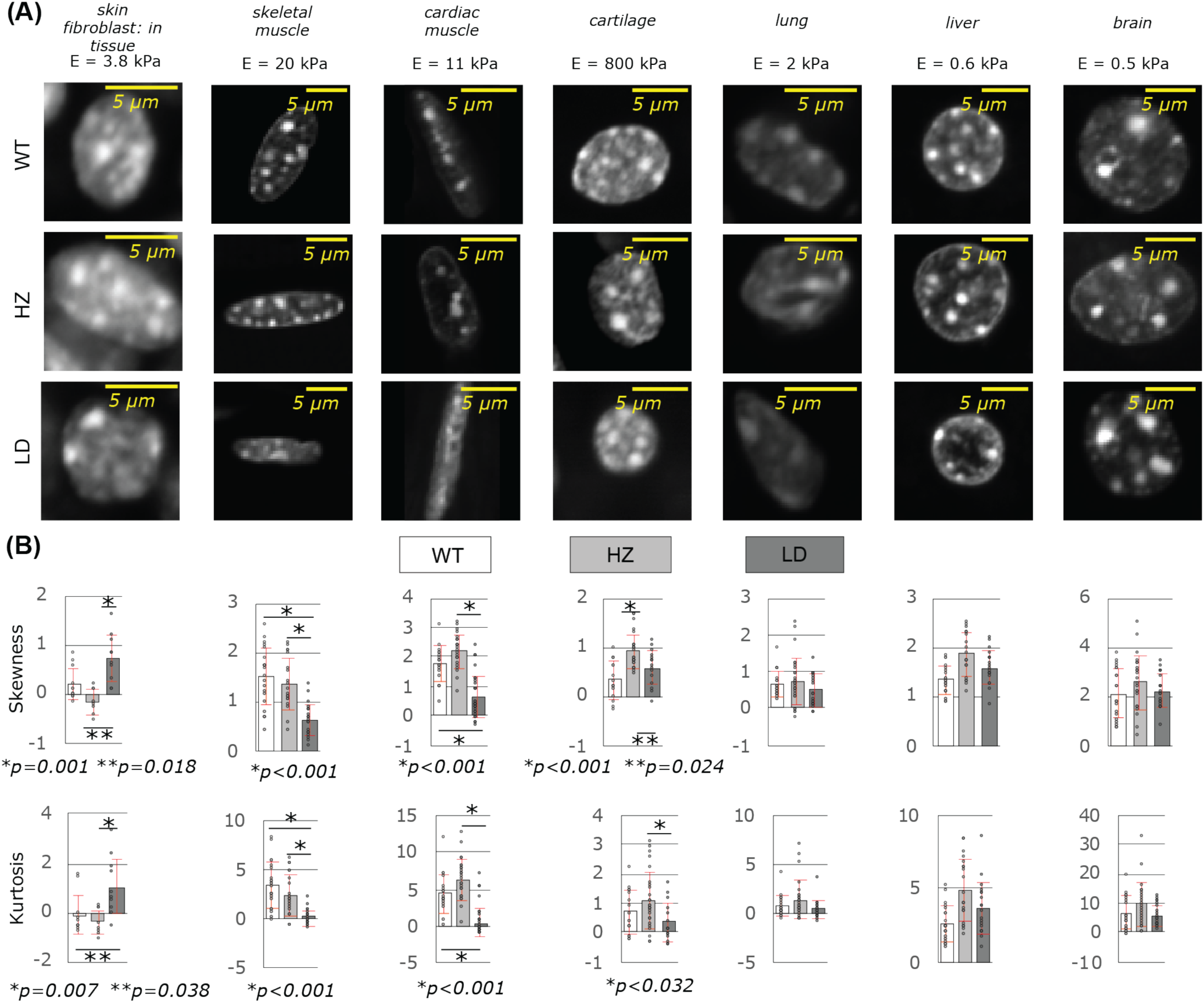
Lamin A/C deficiency disrupts chromatin architecture in mechanically stiff tissues. **(A)** High-resolution DAPI images of individual nuclei from multiple murine tissues, generated using confocal microscopy. DAPI-dense regions correspond to DNA-rich heterochromatin-like domains, whereas lower-intensity regions correspond to more euchromatin-like domains. In LD nuclei from mechanically stiff tissues, including skeletal muscle (tibialis anterior), heart, skin, and partially in cartilage, the distinction between DAPI-dense and DAPI-light regions became less pronounced. This phenotype was not visually evident in softer tissues such as lung, liver, and brain. **(B)** Quantification of intranuclear DAPI intensity distributions using skewness and kurtosis. Higher values indicate stronger spatial segregation between dense and less dense chromatin regions, whereas lower values indicate a more homogeneous DNA-density landscape. Lamin A/C deficiency reduced chromatin segregation in mechanically stiff tissues but not in softer tissues. Data are based on >20 nuclei per group per tissue from at least three animals.

In DAPI-stained nuclei (Figure 3A), densely stained regions correspond to DNA-rich heterochromatin, whereas regions with lower DAPI intensity correspond more closely to euchromatin. In WT nuclei, these regions were visually distinct in several tissues, producing a clear separation between dense chromatin foci and less dense intranuclear regions. In Lamin A/C-deficient nuclei, this distinction became less pronounced in mechanically stiff tissues. Skeletal muscle, heart, skin, and, to a lesser extent, cartilage showed a more diffuse DAPI intensity pattern, suggesting reduced spatial segregation between heterochromatin-rich and euchromatin-rich regions. In contrast, softer tissues such as lung, liver, and brain did not show the same visually evident disruption of chromatin organization.

To quantify these changes, we measured the skewness and kurtosis of intranuclear DAPI intensity distributions. These parameters provide a quantitative readout of chromatin segregation: higher values indicate a more heterogeneous intensity distribution with clearer separation between dense and less dense chromatin regions, whereas lower values indicate a more homogeneous distribution. Consistent with the visual observations, Lamin A/C-deficient nuclei from mechanically stiff tissues showed reduced skewness and kurtosis compared to WT and HZ nuclei, indicating reduced chromatin segregation (Figure 3B). This effect was most evident in skeletal muscle, heart, and skin, and was less apparent or absent in softer tissues.

These findings suggest that chromatin architecture is a more sensitive *in vivo* phenotype of Lamin A/C deficiency than nuclear shape. While intact tissue architecture appears to preserve the gross nuclear outline, it does not fully preserve the internal organization of chromatin. The tissue dependence of this phenotype is also important: Lamin A/C deficiency preferentially disrupted chromatin architecture in mechanically stiff tissues, where Lamin A/C is expected to play a larger role in nuclear mechanical integrity. Thus, the effect of Lamin A/C deficiency *in vivo* is not simply a generalized nuclear deformation phenotype, but a tissue-context-dependent disruption of intranuclear chromatin organization.

### In vivo multiscale mechanics reveals reduced effective nuclear stiffness after Lamin A/C loss

Having found that Lamin A/C deficiency disrupts chromatin architecture in mechanically stiff tissues, we next asked whether this architectural change was accompanied by altered nuclear mechanics *in vivo*. This question is technically challenging because the nucleus must be mechanically loaded and imaged inside an intact living tissue, while preserving the native tissue architecture and physiological force transmission. To achieve this, we used our previously established live-animal deformation microscopy approach^31^, combining controlled hindlimb muscle stimulation with high-resolution confocal imaging of skeletal muscle nuclei and the surrounding tissue *in vivo* (Figure 4A).

**Figure 4.**
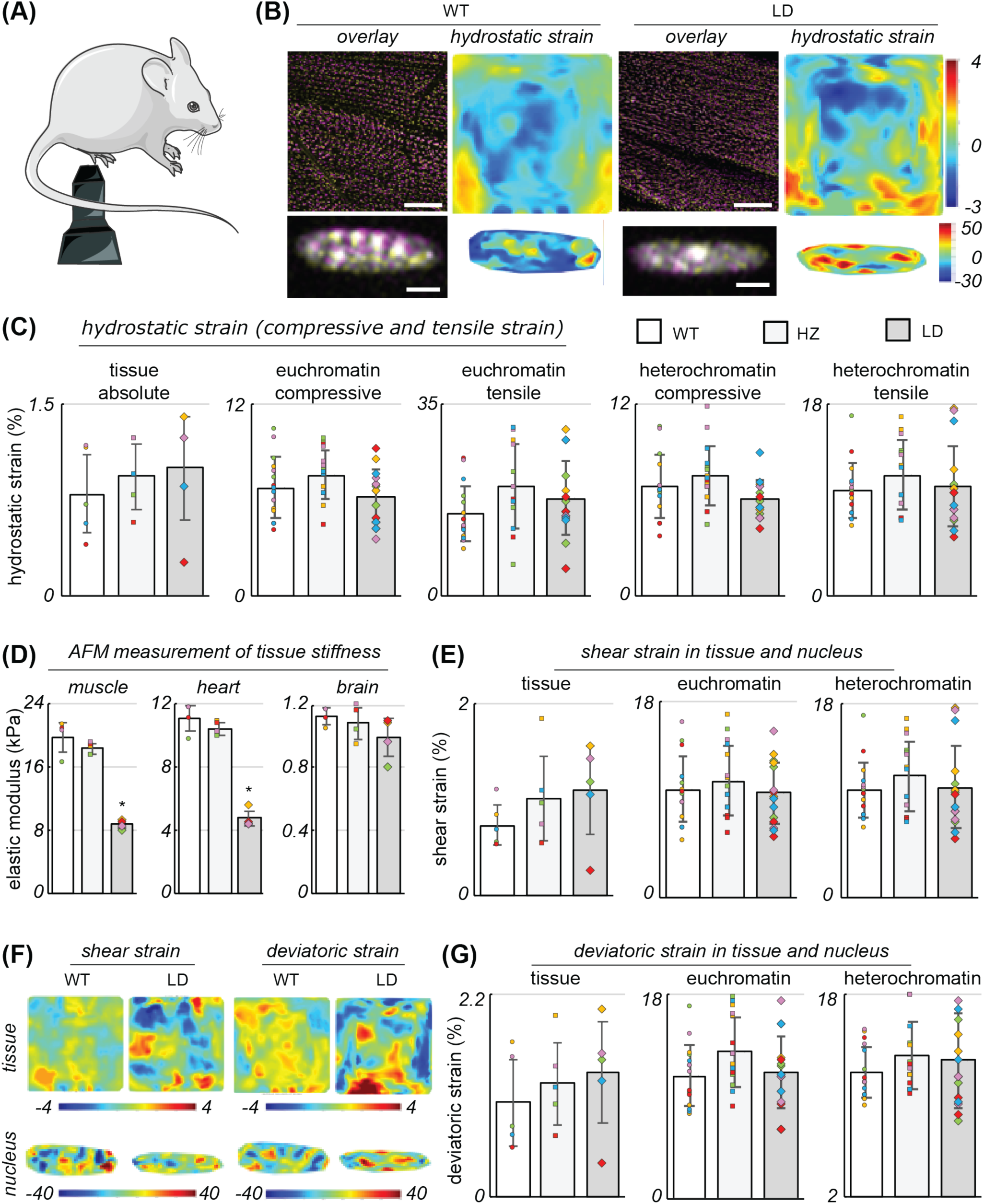
Live in vivo confocal deformation microscopy reveals reduced effective nuclear stiffness after Lamin A/C loss. **(A)** Experimental setup for live-animal multiscale mechanical characterization of skeletal muscle (gastrocnemius). Controlled hindlimb stimulation was used to mechanically load muscle in vivo while high-resolution confocal microscopy captured deformation of the tissue matrix and individual nuclei. **(B)** Representative undeformed and deformed images of the tissue matrix and nucleus, with corresponding hydrostatic strain maps calculated by deformation microscopy. **(C)** Tissue-scale hydrostatic strain was controlled across WT, HZ, and LD groups by adjusting stimulation conditions. Nuclear strain was quantified separately in euchromatin-rich and heterochromatin-rich regions. Average tissue and nuclear strains were comparable across genotypes under matched deformation. **(D)** Young’s modulus of skeletal muscle (tibialis anterior), heart, and brain measured by atomic force microscopy. Lamin A/C deficiency reduced stiffness in skeletal muscle and heart, but not in brain. p < 0.001; n = 4 animals per group. **(E)** Tissue-scale shear strain scaled with intranuclear shear strain across all experimental groups. **(F)** Representative shear and deviatoric strain maps in tissue and nuclei from WT, HZ, and LD mice. **(G)** Tissue-scale deviatoric strain scaled with intranuclear deviatoric strain across groups. Together, these data show that Lamin A/C loss does not simply amplify average nuclear strain under matched tissue deformation. Instead, when combined with the lower tissue stiffness in LD muscle, the strain-scaling analysis suggests reduced effective nuclear stiffness in vivo. In all plots, same color of markers (circle, square, diamond) correspond to data from the same animal.

In this experimental setup, electrical stimulation of the murine hindlimb induced contraction of the tibialis anterior, while the gastrocnemius muscle on the posterior side of the limb was imaged under passive tensile loading. This allowed us to visualize how tissue-scale deformation is transmitted to individual nuclei inside intact skeletal muscle in a live mouse. Confocal images were acquired before and during deformation, and displacement fields were used to compute spatial strain maps in both the tissue matrix and the nucleus (Figure 4B, Figure S6). Because all measurements were performed *in vivo*, the resulting strain fields reflect the native multiscale mechanical environment of skeletal muscle rather than a simplified ex vivo or monolayer culture system.

We first quantified tissue stiffness by atomic force microscopy and found that Lamin A/C deficiency reduced the stiffness of mechanically active tissues (Figure 4B). Both skeletal muscle and heart showed an approximately two-fold reduction in Young’s modulus in Lamin A/C-deficient mice, whereas brain, a softer tissue, did not show a comparable change (Figure 4D, Figure S5). This tissue-level result is consistent with the tissue-specific chromatin phenotype described above and suggests that Lamin A/C deficiency preferentially alters the mechanics of stiffer tissues.

We then used the *in vivo* deformation measurements to determine how tissue deformation was transmitted to the nucleus. The stimulation conditions were adjusted to produce comparable tissue-level hydrostatic strain across WT, HZ, and LD groups (Figure 4C, Figure S6). Under these matched deformation conditions, average nuclear strain was also similar across genotypes when quantified in both euchromatin-rich and heterochromatin-rich regions. Shear and deviatoric strain in the tissue similarly scaled with shear and deviatoric strain inside the nucleus across all groups (Figure 4E, F, G). Thus, Lamin A/C deficiency did not simply produce a larger average nuclear strain under matched tissue deformation.

However, the interpretation changes when the tissue stiffness measurements are considered together with the *in vivo* strain measurements. Because LD skeletal muscle was substantially softer than WT and HZ muscle, the same tissue-level strain would be generated under lower tissue-level stress. If stress transfer from the tissue to the nucleus scales with the applied tissue stress, then similar nuclear strain under lower stress implies a reduced effective nuclear stiffness in Lamin A/C-deficient muscle. In this sense, Lamin A/C loss does not appear as an amplification of average nuclear strain *in vivo*, but rather as a reduction in the effective stiffness of the nucleus within a softer tissue mechanical environment.

Therefore, live *in vivo* deformation microscopy reveals a more nuanced mechanical phenotype than would be expected from nuclear shape alone. Lamin A/C-deficient nuclei preserve their gross morphology and undergo similar average deformation under matched tissue strain, yet the combined tissue stiffness and strain-scaling analysis indicates that their effective nuclear stiffness is reduced. This finding is consistent with previous *in vitro* studies showing that Lamin A/C deficiency softens the nucleus, but here we confirm this mechanical consequence directly in a living tissue environment for the first time. Together, these findings support the idea that Lamin A/C maintains nuclear mechanical homeostasis in intact skeletal muscle, even when the nuclear outline appears largely preserved.

### Lamin A/C deficiency disrupts the load-sharing behavior of euchromatin and heterochromatin in vivo

Although the average nuclear strain was similar across WT, HZ, and LD groups under matched tissue deformation, the nucleus is not a mechanically uniform material. Instead, it contains mechanically and structurally distinct chromatin domains, including more open euchromatin-rich regions and denser heterochromatin-rich regions. We therefore asked whether Lamin A/C deficiency alters how mechanical deformation is distributed inside the nucleus, even when the average nuclear strain appears unchanged.

Using the high-resolution *in vivo* confocal deformation images, we segmented intranuclear regions into euchromatin-rich and heterochromatin-rich domains based on DAPI intensity, as established in our previous work^24^. We then quantified the spatial distribution of hydrostatic strain within each chromatin domain (Figure 5). In WT nuclei, euchromatin and heterochromatin showed a balanced but heterogeneous strain distribution. Euchromatin carried comparable fractions of tensile and compressive strain, whereas heterochromatin showed a modestly greater contribution to tensile strain. This pattern is consistent with a mechanically integrated nucleus in which different chromatin domains participate in load sharing during tissue-scale deformation^26,33^.

**Figure 5.**
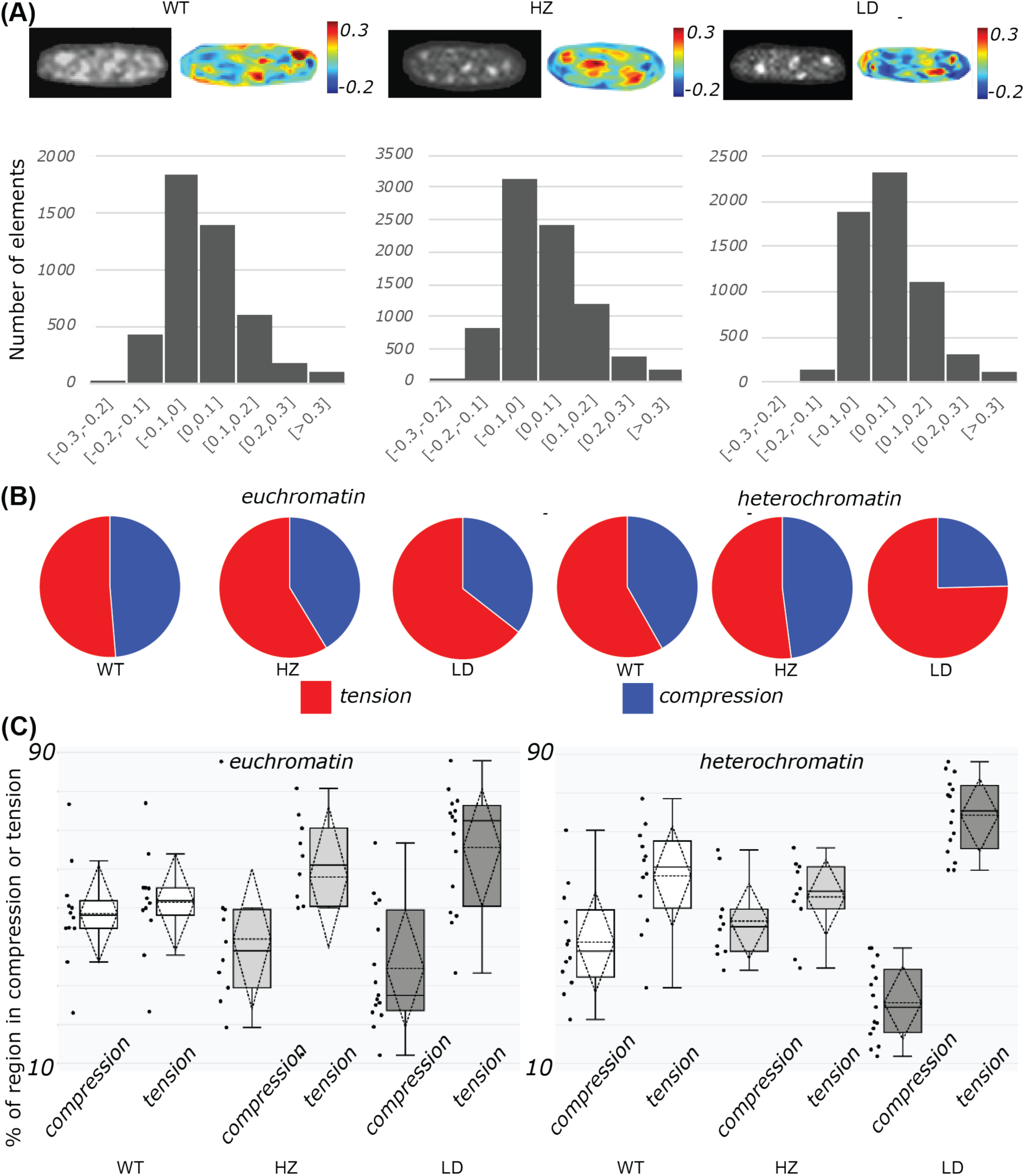
Lamin A/C deficiency disrupts the load-sharing behavior of euchromatin and heterochromatin in vivo. **(A)** Representative hydrostatic strain maps of skeletal muscle (gastrocnemius) nuclei from WT, HZ, and LD mice acquired during live in vivo deformation microscopy. Histograms of nuclear strain values show that tensile and compressive strains are heterogeneously distributed within the nucleus. **(B)** Representative quantification of tensile and compressive strain fractions within euchromatin-rich and heterochromatin-rich domains. **(C)** Group-level analysis showing that Lamin A/C deficiency alters the relative contribution of chromatin domains to intranuclear load sharing. WT nuclei showed a balanced distribution of tensile and compressive strain between euchromatin and heterochromatin, whereas LD nuclei showed increased tensile strain, particularly within heterochromatin-rich regions. HZ nuclei showed altered strain partitioning relative to WT, suggesting that partial Lamin A/C disruption is sufficient to remodel intranuclear mechanics. These data indicate that Lamin A/C maintains the mechanical organization of chromatin domains during tissue-scale deformation.

This load-sharing behavior was disrupted by Lamin A/C deficiency (Figure 5C). In LD nuclei, a larger fraction of euchromatin-rich regions experienced tensile strain, and the effect was even more pronounced in heterochromatin-rich regions, where tensile strain became the dominant mechanical mode. Thus, although the mean nuclear strain did not substantially differ across genotypes, the internal mechanical organization of the nucleus was markedly altered in the absence of Lamin A/C. This suggests that heterochromatin, in particular, loses part of its normal load-bearing behavior when Lamin A/C is disrupted suggesting a loss of heterochromatin stiffness.

Importantly, HZ nuclei also showed evidence of altered intranuclear mechanics (Figure 5C). The HZ group did not simply behave as an intermediate state between WT and LD. Instead, HZ nuclei showed increased tensile loading in euchromatin-rich regions, suggesting that partial Lamin A/C disruption is sufficient to remodel how chromatin domains participate in nuclear deformation. This observation is important because it indicates that chromatin mechanics are already altered in HZ nuclei, even though gross nuclear shape and much of the transcriptomic profile remain closer to WT.

Together, these data show that Lamin A/C deficiency alters the internal mechanical behavior of chromatin before or beyond what is visible from nuclear shape alone. In both HZ and LD nuclei, the normal mechanical partitioning between euchromatin and heterochromatin is disturbed, with LD showing the strongest loss of heterochromatin load sharing. This provides a mechanical basis for the chromatin-architecture changes observed in stiff tissues and led us to ask whether the spatial organization of repressive chromatin marks is also disrupted by partial and complete Lamin A/C loss.

### Partial and complete Lamin A/C disruption uncouple H3K9me3 from DAPI-dense heterochromatin

The altered intranuclear strain distribution in HZ and LD nuclei suggested that chromatin domains may not only deform differently, but may also lose part of their normal epigenetic organization. Because heterochromatin is mechanically and spatially connected to the nuclear lamina through lamina-associated domains, we next asked whether Lamin A/C disruption changes the localization of repressive chromatin marks inside skeletal muscle nuclei. We focused on H3K9me3, a canonical heterochromatin-associated modification, and compared its spatial distribution with DAPI-dense chromatin using Airyscan super-resolution microscopy (Figure 6).

**Figure 6.**
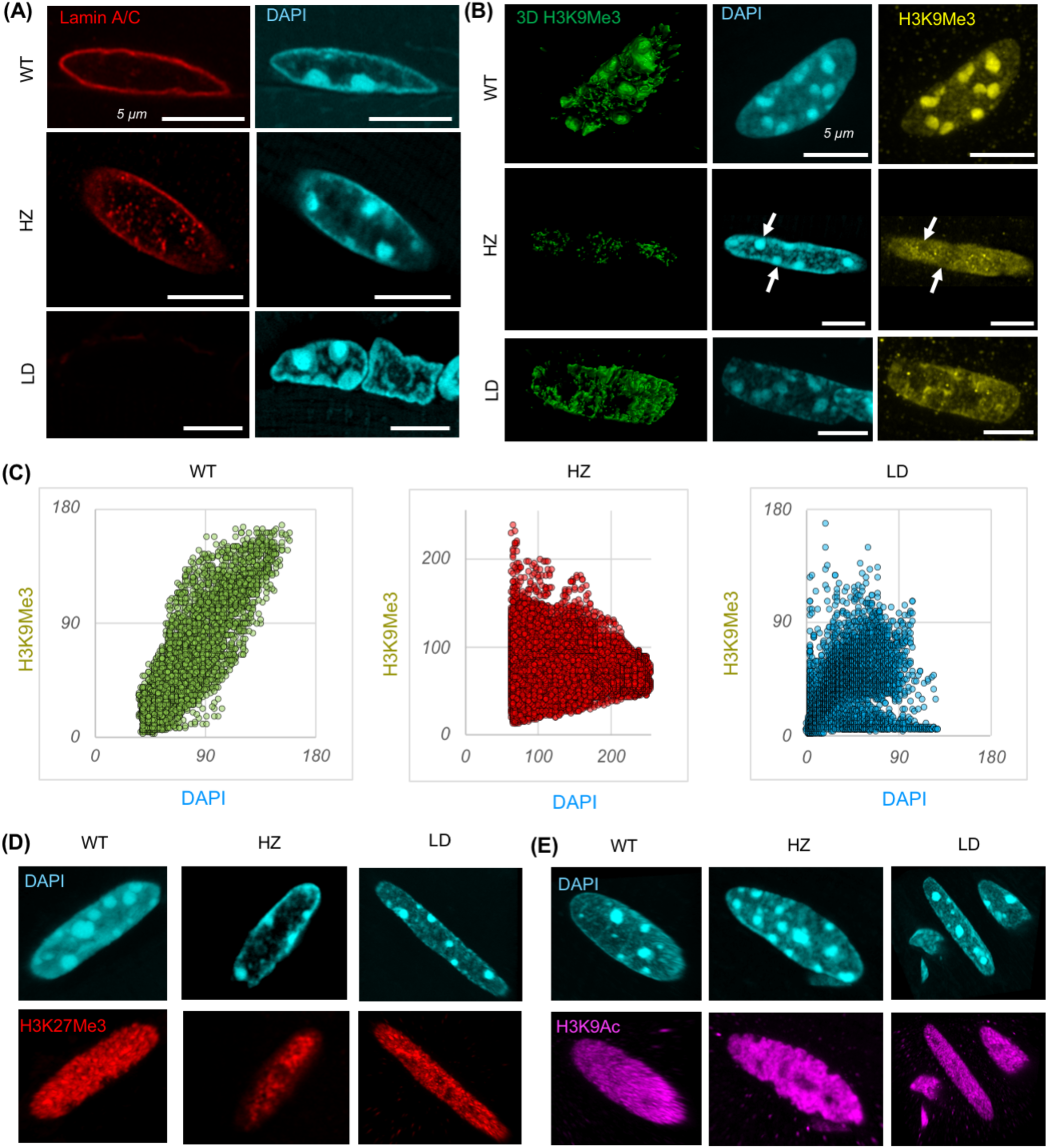
Partial and complete Lamin A/C disruption uncouple H3K9me3 from DAPI-dense heterochromatin. Airyscan super-resolution imaging was used to examine Lamin A/C localization and chromatin-mark organization in skeletal muscle (tibialis anterior) nuclei from WT, HZ, and LD mice. **(A)** Lamin A/C localized primarily to the nuclear envelope in WT nuclei. LD nuclei lacked detectable Lamin A/C signal while retaining Lamin B at the nuclear envelope. HZ nuclei retained peripheral Lamin A/C but also showed Lamin A/C signal within the nuclear interior. **(B)** In WT nuclei, H3K9me3 was spatially coupled to DAPI-dense chromatin domains. In LD nuclei, H3K9me3 appeared more diffuse and was no longer clearly enriched at DAPI-dense regions. HZ nuclei also showed disrupted H3K9me3 organization, with diffuse signal and frequent displacement from DAPI-dense foci. **(C)** Scatterplot of DAPI intensity and H3K9Me3 intensity (after background subtraction) for the images in (B). **(D)** H3K9ac showed a more uniform intranuclear distribution across genotypes. **(E)** H3K27me3 also appeared broadly diffuse and did not show the same genotype-dependent localization pattern as H3K9me3. Together, these data show that partial and complete Lamin A/C disruption uncouple H3K9me3 from DAPI-dense heterochromatin.

In WT skeletal muscle nuclei, Lamin A/C was enriched at the nuclear periphery, consistent with its expected localization at the nuclear lamina (Figure 6A). H3K9me3 showed a spatially organized pattern that was coupled to DAPI-dense chromatin regions (Figure 6B). This organization is consistent with a normal heterochromatin architecture in which compact chromatin domains remain associated with repressive epigenetic marking.

In LD nuclei, Lamin A/C signal was absent (Figure 6A), while Lamin B remained detectable at the nuclear envelope (Figure S3). Although DAPI-dense chromatin foci were still visible in these nuclei, H3K9me3 no longer showed the same spatial relationship with these domains (Figure 6B). Instead, the H3K9me3 signal appeared more diffuse throughout the nucleus and was no longer clearly enriched at DAPI-dense chromatin regions. This indicates that complete Lamin A/C loss disrupts the spatial coupling between compact chromatin and a major repressive histone modification.

Importantly, HZ nuclei also showed a disrupted H3K9me3 organization. Although Lamin A/C remained detectable at the nuclear envelope in HZ nuclei, additional Lamin A/C signal was observed within the nuclear interior (Figure 6A). At the same time, H3K9me3 became more diffuse and, in many nuclei, appeared displaced from or excluded from DAPI-dense chromatin foci rather than colocalized with them (Figure 6B). Thus, partial Lamin A/C disruption is sufficient to alter the spatial organization of H3K9me3, even though HZ nuclei retain peripheral Lamin A/C and preserve a largely WT-like gross nuclear shape. These findings are in line with previous in vitro findings^21^ with the same mouse model, where Lamin A/C was found inside the nucleus in the LD mice embryonic fibroblasts *in vitro*.

We quantified the spatial distribution of H3K9me3 intensity with respect to DAPI intensity to further evaluate this phenotype (Figure 6C). Pearson’s correlation coefficient showed that in WT the DAPI and H3K9Me3 was highly correlated (r = 0.86 for the representative scatterplot) in every nucleus. For HZ nuclei, the correlation was very poor (r = -0.13 for the representative scatterplot) for every nuclei. Interestingly, for LD nuclei, we found a range of correlation coefficient values with very poor correlation (r = -0.5 to moderate correlation r = 0.75; r = 0.71 for the representative scatterplot).

This distinction is important because it suggests that Lamin A/C disruption does not simply erase heterochromatin or uniformly decondense the nucleus. Rather, it separates two normally linked features of nuclear organization: DNA compaction and repressive histone marking. DAPI-dense chromatin domains can remain visible, especially in HZ and LD nuclei, while H3K9me3 becomes mislocalized relative to those domains. In contrast, H3K27me3 (Figure 6D) and H3K9ac (Figure 6E) appeared more uniformly distributed across nuclei and did not show the same genotype-dependent relocalization pattern, suggesting that the spatial effect of Lamin A/C disruption is more evident for H3K9me3 than for other epigenetic modifier marks equally.

Together, these data provide direct imaging evidence that both partial and complete Lamin A/C disruption uncouple H3K9me3 from DAPI-dense heterochromatin in skeletal muscle nuclei. This finding is critical because it shows that HZ nuclei are not simply normal nuclei with reduced Lamin A/C dosage. Rather, they already carry a measurable epigenetic-architectural defect, despite retaining gross nuclear morphology and partial lamina structure. This provided the basis for asking whether partial Lamin A/C disruption also changes chromatin accessibility at the genome-wide level.

### Partial Lamin A/C loss is associated with increased chromatin accessibility in skeletal muscle

The super-resolution imaging data showed that H3K9me3 becomes spatially uncoupled from DAPI-dense heterochromatin in both HZ and LD skeletal muscle nuclei. This suggested that Lamin A/C disruption may alter chromatin regulation beyond what is visible from DNA density alone. To obtain a genome-wide view of this possibility, we performed ATAC-seq profiling in quadriceps muscle from WT, HZ, and LD mice (Figure 7). Because this experiment was performed with one animal per genotype, we interpreted these data as a supportive and hypothesis-generating extension of the imaging results rather than as a standalone definitive measure of chromatin accessibility.

**Figure 7.**
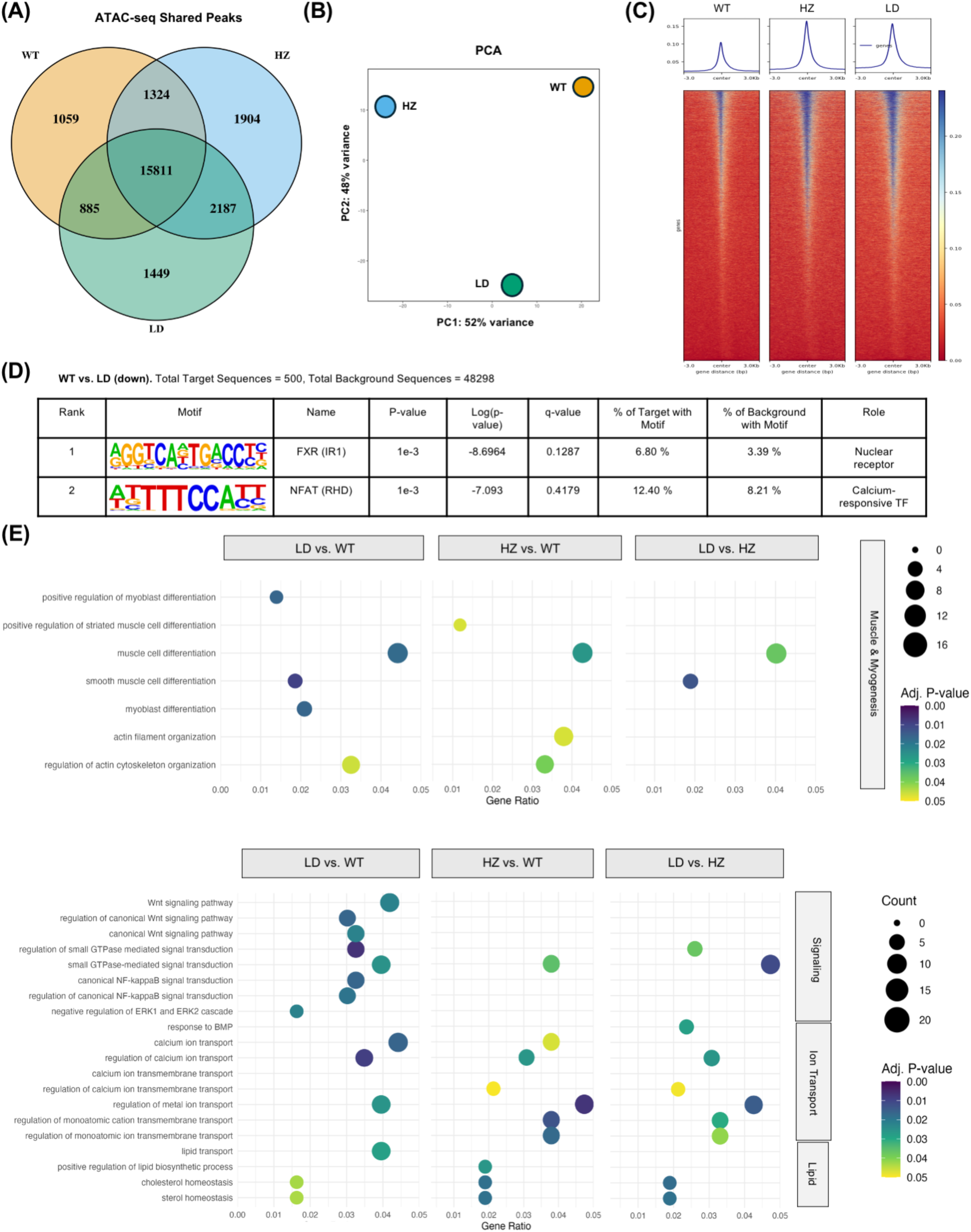
Exploratory ATAC-seq suggests increased chromatin accessibility in HZ skeletal muscle. ATAC-seq was performed (n=1 mouse/ group) on quadriceps muscle from WT, HZ, and LD mice to obtain a genome-wide view of chromatin accessibility. Because this analysis was performed with one animal per genotype, the data were interpreted as supportive and hypothesis-generating rather than as a standalone definitive measure of chromatin accessibility. **(A)** Venn diagram showing shared and genotype-specific accessible chromatin peaks. Most peaks were shared across all three groups, while HZ showed the largest number of unique accessible regions. **(B)** Principal component analysis of ATAC-seq profiles. Although clustering cannot be interpreted definitively due to the limited sample number, HZ occupied a distinct position relative to WT and LD. **(C)** ATAC-seq signal enrichment around transcription start sites showed higher accessibility in HZ compared to WT and LD, suggesting increased regulatory-region accessibility after partial Lamin A/C loss. **(D)** Motif enrichment analysis of differentially accessible regions suggested altered accessibility at sites associated with transcription factors involved in calcium signaling and nuclear receptor pathways. **(E)** Gene Ontology biological process analysis of differentially accessible regions identified pathways related to muscle structure, myogenesis, cellular signaling, ion transport, and lipid metabolism. Together, these exploratory data are consistent with the imaging evidence that HZ muscle undergoes chromatin-level remodeling despite preserved gross nuclear morphology.

Most accessible chromatin regions were shared across all three genotypes, indicating that a large fraction of the skeletal muscle accessibility landscape was preserved despite Lamin A/C disruption (Figure 7A). At the same time, each genotype also contained a set of unique peaks, with HZ showing the largest number of unique accessible regions among the three groups. Principal Component Analysis (PCA) did not show clear genotype clustering, which is expected given the limited sample number, but the HZ sample did not simply overlap with either WT or LD. Instead, it occupied a distinct position in the accessibility landscape (Figure 7B).

Consistent with this observation, ATAC-seq signal around transcription start sites was higher in HZ muscle compared to WT and LD (Figure 7C). This pattern suggests that partial Lamin A/C loss may be associated with increased chromatin accessibility near regulatory regions. Importantly, this result agrees with the preceding imaging data (Figure 6B), where HZ nuclei already showed altered Lamin A/C localization and disrupted H3K9me3 organization despite maintaining gross nuclear shape. Thus, the ATAC-seq data support the idea that HZ muscle is not same as WT, but undergoes a chromatin-level adaptation that is not apparent from nuclear morphology alone.

Gene ontology analysis of differentially accessible regions further suggested that chromatin remodeling in HZ and LD muscle may involve pathways related to muscle structure, myogenesis, cellular signaling, ion transport, and lipid metabolism (Figure 7E). In particular, WT versus HZ comparisons included terms related to striated muscle differentiation and actin filament organization, whereas WT versus LD comparisons showed strong enrichment of muscle-related pathways. Motif analysis also suggested altered accessibility at sites associated with transcription factors involved in calcium signaling and nuclear receptor pathways (Figure 7D). These observations are consistent with the idea that Lamin A/C disruption affects regulatory regions connected to muscle function and mechanosensitive signaling.

Overall, the ATAC-seq data provide a genome-wide view that is consistent with the imaging-based chromatin phenotype. They suggest that partial Lamin A/C loss is associated with increased chromatin accessibility in skeletal muscle, especially near transcription start sites. However, because of the limited sample number, we use these data primarily to support and extend the direct imaging observations rather than to define the central mechanism by themselves. Together with the H3K9me3 redistribution data, these findings suggest that HZ nuclei undergo chromatin remodeling before the emergence of any phenotype at the transcriptomic or proteomic level. This led us to ask whether these chromatin-level changes are accompanied by corresponding changes in gene expression.

### Complete Lamin A/C loss disrupts muscle transcription, whereas partial loss preserves a near-WT transcriptomic state

Because the ATAC-seq data suggested chromatin-accessibility changes in HZ muscle, we next asked whether these changes were accompanied by broad transcriptional disruption. To address this, we performed bulk RNA-seq on quadriceps muscle from WT, HZ, and LD mice (n = 4 animals/ group) (Figure 8). This analysis allowed us to determine whether the chromatin remodeling observed in HZ and LD nuclei was associated with corresponding changes in gene expression.

**Figure 8.**
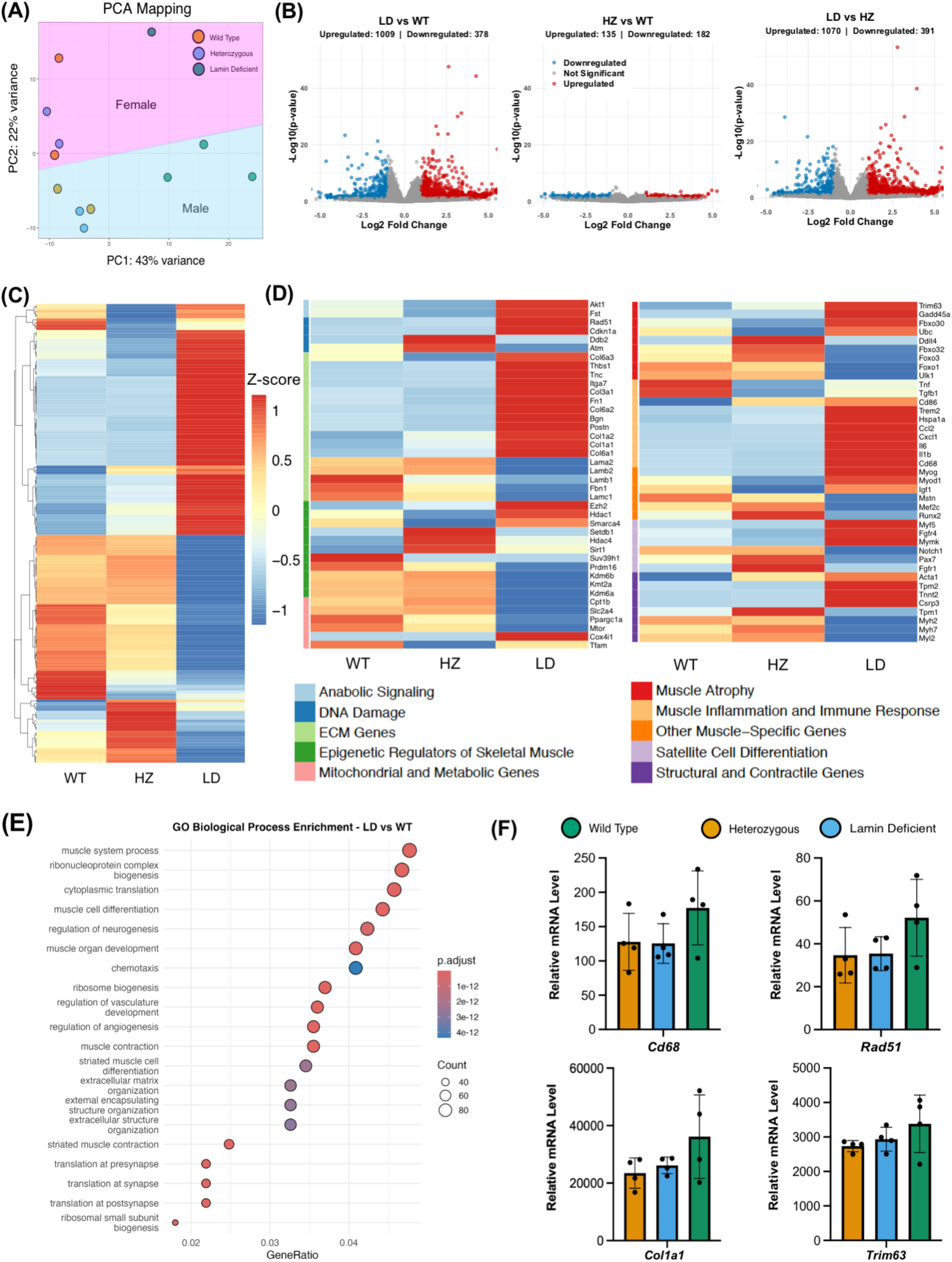
Complete Lamin A/C loss disrupts muscle transcription, whereas partial loss preserves a near-WT transcriptomic state. RNA-seq was performed on quadriceps muscle from WT, HZ, and LD mice, with n = 4 animals per group. **(A)** Principal component analysis showed clear separation of LD samples from WT and HZ along PC1, indicating that complete Lamin A/C loss is the major driver of transcriptomic variation. PC2 separated samples by sex, indicating a secondary contribution of sex to gene-expression variation. **(B)** Volcano plots of pairwise comparisons showed extensive differential gene expression in LD versus WT and LD versus HZ comparisons, whereas HZ versus WT showed fewer changes. **(C)** Z-score heatmap of normalized gene expression showing broad transcriptomic divergence of LD muscle, while HZ and WT retained more similar expression patterns. **(D)** Heatmap of genes grouped by functional categories showing marked disruption of muscle, stress, extracellular matrix, inflammatory, and chromatin-related programs in LD muscle. HZ muscle remained closer to WT across many categories but showed selective changes in specific regulatory genes. **(E)** Gene Ontology biological process analysis of differentially expressed genes. LD versus WT comparisons were enriched for pathways related to muscle structure, contractile function, extracellular matrix remodeling, and neuromuscular synaptic organization. No significantly enriched pathways were detected in the HZ versus WT comparison. **(F)** Relative mRNA expression of representative genes associated with fibrosis, proteolysis, DNA damage, inflammation, and degeneration. WT: orange, HZ: blue, LD: green. Col1a1, Trim63, Rad51, and Cd68 were elevated in LD muscle but remained closer to WT levels in HZ muscle. These data show that complete Lamin A/C loss produces broad myopathic transcriptional dysregulation, whereas partial loss preserves a near-WT transcriptomic state with selective gene-level adaptation.

PCA revealed that genotype was the major source of transcriptomic variation (Figure 8A). LD samples separated clearly from WT and HZ along PC1, indicating that complete Lamin A/C loss produces a distinct transcriptional state. In contrast, HZ samples clustered closer to WT, suggesting that partial Lamin A/C loss does not lead to the same broad transcriptomic disruption as complete loss. PC2 separated samples by sex, indicating that sex contributed a secondary source of variation, but did not override the dominant effect of complete Lamin A/C deficiency.

Pairwise differential expression analysis further supported this pattern (Figure 8B). LD muscle showed extensive gene-expression changes compared to both WT and HZ, whereas HZ muscle showed a much smaller transcriptional shift relative to WT. Heatmap analysis of normalized gene expression also showed that HZ and WT shared similar overall expression patterns across many functional categories, while LD muscle diverged more strongly (Figure 8C, D). Thus, the transcriptomic phenotype was not simply proportional to Lamin A/C level. Instead, partial Lamin A/C loss was associated with chromatin remodeling but relative preservation of the global muscle transcriptome.

Despite this overall preservation, HZ muscle was not transcriptionally identical to WT (Figure 8D). Several genes involved in epigenetic regulation, muscle transcription, and cellular stress showed selective changes in HZ. For example, *Hdac4*, *Sirt1*, *Mef2c*, and *Fbxo32* showed higher expression in HZ compared to WT and LD, whereas genes such as *Kdm4a* and *Smarca4* showed lower expression. Other stress- and chromatin-associated genes, including *Setdb1* and *Ddit4*, also showed HZ-specific changes. These gene-level differences suggest that the near-WT transcriptomic state in HZ is not passive, but may reflect a selective compensatory response to partial Lamin A/C disruption.

Gene ontology analysis emphasized the difference between partial and complete Lamin A/C loss (Figure 8E). LD versus WT differentially expressed genes were enriched for pathways related to muscle structure, contractile function, extracellular matrix remodeling, and neuromuscular synaptic organization. These pathways are consistent with the known muscle pathology associated with Lamin A/C deficiency. In contrast, the HZ versus WT comparison did not yield significantly enriched pathways, suggesting that partial Lamin A/C loss does not produce the same pathway-level transcriptional failure.

We further examined representative genes associated with fibrosis, proteolysis, DNA damage, inflammation, and muscle degeneration (Figure 8F). Genes such as *Col1a1*, *Trim63*, *Rad51*, and *Cd68* were elevated in LD muscle but remained closer to WT levels in HZ muscle. This pattern supports the idea that HZ muscle preserves many transcriptional features of WT tissue, whereas LD muscle enters a broader pathological state involving extracellular matrix remodeling, inflammatory activation, stress response, and degeneration-associated pathways.

Together, these RNA-seq data reveal a nonlinear transcriptional response to Lamin A/C level in skeletal muscle *in vivo*. Complete Lamin A/C loss causes broad disruption of the skeletal muscle transcriptome, while partial loss preserves a near-WT expression landscape despite clear evidence of altered intranuclear mechanics, H3K9me3 organization, and chromatin accessibility. This suggests that HZ muscle engages compensatory mechanisms that buffer gene expression against partial disruption of the nuclear lamina. The next question, therefore, was whether the chromatin-accessibility and transcriptomic data could identify candidate pathways that mediate this compensation.

### Integrated chromatin-accessibility and transcriptomic analysis suggests an HDAC2-dependent compensatory mechanism in HZ muscle

The preceding data showed an apparent paradox in HZ muscle. Partial Lamin A/C disruption altered intranuclear mechanics, uncoupled H3K9me3 from DAPI-dense heterochromatin, and was associated with increased chromatin accessibility, yet the global transcriptomic state remained close to WT. We therefore asked whether integrating chromatin-accessibility and RNA-seq data could identify candidate regulatory mechanisms that help preserve transcriptional homeostasis in HZ muscle (Figure 9, Figure S7).

**Figure 9.**
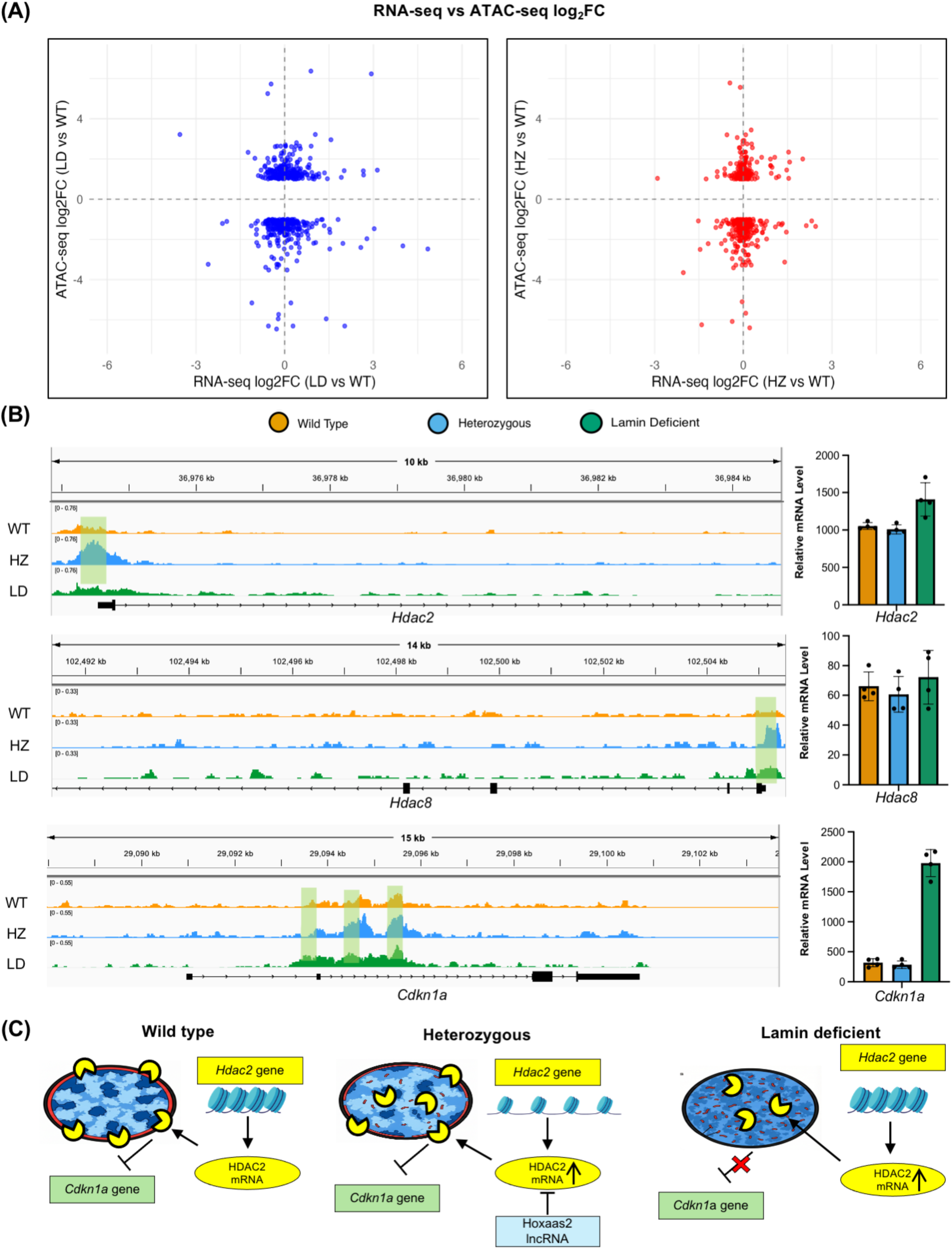
Integrated chromatin-accessibility and transcriptomic analysis suggests an HDAC2-dependent compensatory mechanism in HZ muscle. **(A)** Correlation analysis comparing promoter-associated differentially accessible regions with differentially expressed genes in LD versus WT and HZ versus WT comparisons. Data points were distributed across all four quadrants, indicating weak global coupling between promoter accessibility and gene expression. This suggests that Lamin A/C-dependent chromatin remodeling does not translate into transcriptional change through a simple linear relationship. **(B)** ATAC-seq tracks and relative mRNA expression for candidate regulatory genes. The Hdac2 locus showed increased chromatin accessibility in HZ muscle, while Hdac2 transcript levels remained close to WT. Cdkn1a expression was elevated in LD muscle but remained near WT levels in HZ, consistent with preserved repression of a stress- and cell-cycle-associated gene after partial Lamin A/C disruption. Green-highlighted regions indicate accessible chromatin peaks. **(C)** Proposed model. In WT muscle nuclei, Lamin A/C supports lamina-associated heterochromatin organization and helps maintain repressive chromatin regulation. In HZ nuclei, partial Lamin A/C disruption alters intranuclear mechanics and H3K9me3 organization, but residual lamina structure may allow compensatory chromatin accessibility and HDAC2-dependent buffering of gene expression. In LD nuclei, complete Lamin A/C loss disrupts lamina-associated repression, uncouples H3K9me3 from compact chromatin, and activates Cdkn1a and broader pathological gene programs. This model positions HDAC2 as a candidate mechanosensitive regulatory node linking Lamin A/C-dependent nuclear architecture to transcriptional homeostasis. The HDAC2 mRNA level may be kept under control in the HZ group by several candidates. One of them may be long non-coding RNA Hoxaas2, whose level was higher in HZ compared to WT and HZ, as discovered from transcriptomics data.

We first compared differentially accessible regions with differentially expressed genes across genotypes. This analysis did not show a strong global correlation between promoter accessibility and gene expression (Figure 9A). Data points were distributed across all four quadrants in both LD versus WT and HZ versus WT comparisons, indicating that increased accessibility at a given region did not necessarily correspond to increased expression of the nearby gene. This weak global coupling is important because it suggests that Lamin A/C-dependent chromatin remodeling does not translate into gene expression of openness of chromatin is equivalent to transcriptionally active. Instead, the transcriptional consequences of altered chromatin accessibility appear to be locus-specific and likely shaped by additional regulatory mechanisms. We next examined genes and pathways that could connect Lamin A/C disruption, chromatin accessibility, and transcriptional buffering. Class I histone deacetylases have been implicated in gene repression at the nuclear lamina, including HDAC1- and HDAC3-dependent regulation of lamina-associated chromatin domains^34,35^. HDAC2 is particularly relevant here because Lamin A/C directly interacts with HDAC2 and influences HDAC2 recruitment to the *CDKN1A/p21* promoter^36,37^; disruption of this Lamin A/C-HDAC2 interaction in progeroid cells is associated with impaired p21 repression during stress recovery. In our integrated analysis, the *Hdac2* locus showed increased chromatin accessibility in HZ muscle compared to WT and LD, whereas *Hdac2* mRNA levels remained close to WT (Figure 9B). This pattern is consistent with the broader HZ phenotype: chromatin accessibility is remodeled, but transcript levels are maintained within a near-normal range.

The downstream behavior of Cdkn1a, which encodes p21, further supported this interpretation. Cdkn1a is normally restrained by HDAC2-dependent repression, and its activation is associated with cell-cycle arrest, senescence-like programs, and muscle pathology. In LD muscle, Cdkn1a expression was increased, consistent with failure of Lamin A/C-associated repression. In contrast, HZ muscle maintained Cdkn1a expression closer to WT levels. Thus, despite altered chromatin architecture and increased accessibility, HZ muscle appears to preserve repression of a key stress- and cell-cycle-associated gene that becomes activated after complete Lamin A/C loss.

Based on these observations, we propose that HZ muscle engages a compensatory mechano-epigenetic mechanism centered, at least in part, on HDAC2. In WT nuclei, Lamin A/C supports normal heterochromatin organization and helps maintain repressive chromatin regulation at the nuclear periphery. In HZ nuclei, partial Lamin A/C disruption alters chromatin mechanics and H3K9me3 organization, but residual lamina structure may allow the nucleus to use increased chromatin accessibility as an adaptive state. This may preserve HDAC2-dependent repression of Cdkn1a and other stress-associated genes, thereby maintaining a transcriptomic profile closer to WT. In LD nuclei, this compensation appears to fail - loss of Lamin A/C disrupts lamina-associated organization more completely, H3K9me3 becomes spatially uncoupled from compact chromatin, and Cdkn1a and broader pathological gene programs are activated.

This model (Figure 9C) does not imply that HDAC2 alone explains all transcriptional buffering in HZ muscle. Rather, HDAC2 emerges from the integrated analysis as a candidate mechanosensitive regulatory node that links Lamin A/C-dependent nuclear architecture with gene-expression homeostasis. Other regulatory layers, including additional chromatin modifiers, transcription factors, and non-coding RNAs (such as Hoxaas2), may also contribute. Nevertheless, the HDAC2-Cdkn1a axis provides a plausible locus-specific example of how partial Lamin A/C disruption can remodel chromatin without producing the broad transcriptional failure observed in LD muscle.

Together, these integrated data support a nonlinear model of Lamin A/C dosage in skeletal muscle. Partial Lamin A/C disruption produces measurable mechanical and epigenetic remodeling, but this remodeling is accompanied by transcriptional buffering that preserves a near-WT gene-expression state. Complete Lamin A/C loss exceeds this compensatory capacity, leading to failure of repressive chromatin regulation and broad myopathic transcriptional disruption. Thus, Lamin A/C does not simply maintain nuclear shape *in vivo*; it coordinates nuclear mechanics, chromatin organization, and transcriptional homeostasis in skeletal muscle.

## DISCUSSION

Lamin A/C and its associated proteins have been investigated for more than two decades, yet the physiological role of Lamin A/C in maintaining chromatin architecture, intranuclear mechanics, and gene regulation in intact tissues remains incompletely understood. Much of the foundational knowledge in the field has come from cultured cells, where Lamin A/C deficiency or mutation is associated with abnormal nuclear shape, reduced nuclear stiffness, altered chromatin organization, and disrupted gene expression. These studies have been essential for establishing Lamin A/C as a structural and regulatory component of the nucleus. However, cultured cells, particularly cells grown as two-dimensional monolayers on rigid substrates, do not fully reproduce the mechanical and architectural constraints experienced by nuclei inside tissues. This distinction is especially important for mechanobiology, where cell shape, cytoskeletal organization, extracellular matrix mechanics, and tissue-level force transmission can strongly influence nuclear behavior.

In this study, we investigated Lamin A/C function in intact tissues and in live mouse skeletal muscle. The central finding is that Lamin A/C deficiency does not primarily appear *in vivo* as a generalized collapse of nuclear shape. Instead, it is more sensitive and physiologically relevant phenotype is an internal reorganization of the nucleus: altered chromatin architecture, reduced effective nuclear stiffness, disrupted intranuclear load sharing between euchromatin and heterochromatin, redistribution of H3K9me3, and altered transcriptional homeostasis in skeletal muscle. This distinction is important because it shifts the interpretation of Lamin A/C function from a mostly nuclear-shape-centric model to a mechano-epigenetic model in which Lamin A/C coordinates the mechanical and chromatin-regulatory state of the nucleus in a tissue-context-dependent manner.

One of the most unexpected findings was that nuclear shape was largely preserved across intact tissues despite Lamin A/C deficiency. In 2D monolayer culture, we observed the expected trend: a subpopulation of Lamin A/C-deficient skin-derived and cardiac fibroblasts showed irregular nuclear shape and occasional blebbing. This is consistent with prior *in vitro* studies showing that Lamin A/C contributes to nuclear stiffness and nuclear morphology^6^. However, the same dramatic nuclear-shape phenotype was not observed as a dominant feature in intact tissues. Even skeletal muscle and heart, which are among the most clinically affected tissues in laminopathies, showed only modest and statistically non-significant increases in nuclear irregularity. This suggests that native tissue architecture can mechanically buffer nuclear shape. *In vivo*, cells are not free to spread and define their geometry autonomously as they do on a culture dish; rather, they are integrated within extracellular matrix, neighboring cells, and tissue-level structural organization. These multiscale constraints may preserve the gross nuclear outline even when the nuclear lamina itself is compromised.

This finding does not contradict the extensive *in vitro* Lamin A/C literature which states that Lamin A/C deficient nuclei are softer^6^ or stiffness drives the Lamin A/C content over Lamin B^23^. Instead, it places that literature in a more physiological context. Monolayer culture is a powerful reductionist system because it exposes cell-autonomous defects in nuclear mechanics and morphology. However, the same system can also exaggerate or redirect mechanical phenotypes by imposing artificial substrate stiffness, spreading geometry, and cytoskeletal tension. Our data suggest that the nuclear-shape phenotype observed *in vitro* is not necessarily the dominant phenotype *in vivo*. In intact tissues, Lamin A/C deficiency may be less visible from the outside of the nucleus but more consequential inside the nucleus. This is a key conceptual advance of the study: *in vivo* Lamin A/C dysfunction can be mechanically and epigenetically severe even when nuclear shape appears mostly preserved.

Consistent with this idea, we found that chromatin architecture was more sensitive to Lamin A/C deficiency than nuclear shape. DAPI intensity analysis showed reduced segregation between DNA-dense and DNA-light chromatin regions in mechanically stiff tissues, including skeletal muscle, heart, skin, and cartilage, but not in softer tissues such as brain, liver, and lung. This tissue dependence is consistent with the known relationship between Lamin A/C abundance and tissue stiffness^23^. Lamin A/C is more abundant in mechanically stiff tissues, where nuclei are expected to experience greater mechanical load and where the lamina may be particularly important for stabilizing chromatin architecture. Thus, Lamin A/C deficiency does not simply produce a uniform nuclear phenotype across the body. Rather, its effects depend on the mechanical context of the tissue.

The *in vivo* mechanical measurements further support this interpretation. Using deformation microscopy in live animals, we quantified tissue-to-nucleus strain transfer in skeletal muscle while the tissue remained inside the living mouse. This approach preserves the native mechanical environment of muscle, including physiological tissue architecture and multiscale force transmission. Previous *in vitro* studies have shown that Lamin A/C deficiency softens the nucleus and alters nuclear deformation under applied force^6^. Our study confirms this mechanical consequence *in vivo*, but in a more nuanced manner. Under matched tissue-level strain, average nuclear strain was similar across WT, HZ, and LD groups. However, because Lamin A/C-deficient skeletal muscle was substantially softer, similar nuclear strain under lower inferred stress suggests reduced effective nuclear stiffness in LD muscle. Therefore, Lamin A/C loss *in vivo* does not simply manifest as amplified average nuclear strain; instead, it changes the mechanical state of the tissue-nucleus system.

The most revealing mechanical phenotype was not the average nuclear strain, but the distribution of strain inside the nucleus. WT nuclei showed a more balanced load-sharing behavior between euchromatin-rich and heterochromatin-rich regions. In LD nuclei, this balance was disrupted, with heterochromatin-rich regions showing a stronger shift toward tensile strain. HZ nuclei also showed altered intranuclear strain partitioning, particularly in euchromatin-rich regions. These findings suggest that Lamin A/C helps maintain the mechanical role of chromatin domains during tissue-scale deformation. In this framework, heterochromatin is not only a transcriptionally repressive compartment, but also a mechanically relevant nuclear domain. When Lamin A/C is disrupted, heterochromatin appears to lose part of its normal load-bearing behavior. This provides a physical basis for why changes in nuclear mechanics and chromatin regulation may be coupled having implications in chromatin accessibility.

The H3K9me3 imaging data provide an important epigenetic link to this mechanical phenotype. In WT skeletal muscle nuclei, H3K9me3 was spatially coupled to DAPI-dense heterochromatin domains. In LD nuclei, H3K9me3 became diffuse and was no longer clearly enriched at these dense chromatin regions. Importantly, HZ nuclei also showed disrupted H3K9me3 organization, despite retaining peripheral Lamin A/C and a largely preserved gross nuclear shape. This finding suggests that partial Lamin A/C disruption is sufficient to uncouple a major repressive histone mark from compact chromatin. The interpretation of this result requires care because DAPI intensity and H3K9me3 intensity report different aspects of nuclear organization. DAPI skewness and kurtosis reflect DNA-density segregation, whereas H3K9me3 distribution reflects the concentration and spatial heterogeneity of the histone mark itself. The Lamin A/C-H3K9me3 relationship is likely indirect and mediated by lamina-associated chromatin tethers. LAP2β, LBR, PRR14, HP1, and HDAC-associated complexes have all been implicated in peripheral heterochromatin organization^38^, providing plausible routes by which Lamin A/C disruption could redistribute H3K9me3-marked chromatin.

The genome-wide data support this model while also highlighting the need for cautious interpretation. ATAC-seq suggested that HZ muscle has increased chromatin accessibility, including increased signal around transcription start sites and a larger number of unique accessible regions. The ATAC-seq data are consistent with the direct imaging and mechanics results: HZ nuclei are not normal nuclei with half the Lamin A/C dose. They show altered intranuclear mechanics, disrupted H3K9me3 organization, and evidence of increased chromatin accessibility. The strength of the HZ conclusion therefore does not rest only on ATAC-seq; rather, ATAC-seq provides a genome-wide extension of phenotypes already observed by imaging and mechanical analysis.

The RNA-seq data revealed a nonlinear relationship between Lamin A/C dosage and transcriptional outcome. Complete Lamin A/C loss produced broad transcriptional dysregulation in skeletal muscle, including pathways related to muscle structure, contractile function, extracellular matrix remodeling, inflammation, stress response, and degeneration-associated programs. In contrast, HZ muscle remained much closer to WT at the global transcriptomic level, despite showing altered nuclear mechanics and chromatin organization. This divergence suggests that partial Lamin A/C disruption triggers compensatory mechanisms that buffer gene expression, whereas complete Lamin A/C loss exceeds the capacity of this compensation. This is an important point because it argues against a simple dose-dependent model in which HZ is merely an intermediate state between WT and LD. Instead, HZ appears to represent an adaptive state: mechanically and epigenetically altered, but transcriptionally buffered.

Integrated chromatin-accessibility and transcriptomic analysis suggested HDAC2 as one candidate regulatory node in this compensation. Class I HDACs interact with the nuclear lamina, and HDAC2 has been linked to Lamin A/C-dependent repression of cell-cycle arrest genes. In our data, the *Hdac2* locus showed increased accessibility in HZ muscle, while Hdac2 transcript levels remained close to WT. At the same time, Cdkn1a, which encodes p21 and is normally restrained by HDAC2-dependent repression, remained near WT levels in HZ but was elevated in LD. This pattern supports a model in which partial Lamin A/C disruption increases chromatin accessibility but still preserves sufficient lamina-associated regulatory function to maintain repression of stress- and cell-cycle-associated genes. In LD nuclei, this regulatory state fails, leading to Cdkn1a activation and broader pathological transcriptional remodeling.

This model should be interpreted as a working mechanism rather than a completed causal pathway. Our data identify HDAC2 as a plausible mechanosensitive compensatory node, but they do not prove that HDAC2 alone drives transcriptional buffering in HZ muscle. Functional perturbation of HDAC2, direct mapping of HDAC2 binding, and rescue experiments would be required to establish causality. Similarly, ChIP-seq for H3K9me3 and Hi-C or related chromatin-conformation assays would help determine whether H3K9me3 redistribution corresponds to altered lamina-associated domains, long-range chromatin interactions, or specific changes in heterochromatin compartmentalization. These future studies will be important for determining how mechanical changes in the nucleus are converted into durable epigenetic and transcriptional states.

A key strength of this study is the use of intact tissues and live *in vivo* mechanical measurements, but this also introduces limitations. *In vivo* experiments preserve physiological architecture, but they are technically complex and less reductionist than *in vitro* systems. The inferred effective nuclear stiffness depends on mechanical scaling assumptions, including how tissue-level stress is transmitted to the nucleus. While the combined tissue stiffness and strain data strongly support reduced effective nuclear stiffness in Lamin A/C-deficient muscle, direct measurement of nuclear modulus *in vivo* remains technically difficult. Similarly, chromatin segmentation based on DAPI intensity provides a useful readout of DNA-density organization, but it is not equivalent to direct molecular annotation of euchromatin and heterochromatin. The ATAC-seq analysis, as noted above, is exploratory because of limited sample number. We therefore use it to support the imaging-based chromatin phenotype and to generate mechanistic hypotheses, rather than as the primary basis for the central conclusions.

Another limitation is that skeletal muscle is the main tissue in which multiscale nuclear mechanics, chromatin accessibility, RNA-seq, and epigenetic imaging were integrated. Our tissue survey suggests that Lamin A/C-dependent chromatin disruption is more evident in mechanically stiff tissues, but the complete mechanical and molecular cascade was not tested in every tissue. It remains possible that heart, skin, cartilage, and other stiff tissues use overlapping but distinct compensatory mechanisms. The broader principle suggested by this study is that Lamin A/C function depends on tissue mechanical context, but the exact downstream mechanisms may be tissue specific.

The relevance of these findings to aging should also be viewed in a balanced way. Lamin A/C biology is closely connected to progeroid syndromes, nuclear architecture, genome organization, DNA damage responses, senescence-associated pathways, and tissue degeneration^27,28,39^. In support of a connection to physiological aging, cells from aged healthy individuals have been reported to acquire lamin A-dependent nuclear defects resembling those observed in Hutchinson-Gilford progeria syndrome, including altered histone modifications and increased DNA damage^27^. In addition, impaired lamin A processing and prelamin A accumulation have been linked to vascular smooth muscle cell aging, DNA damage, and senescence-like phenotypes^40^. Therefore, the mechanisms described here may be relevant to aging-related decline in mechanically active tissues. However, this study does not directly test physiological aging. The mice used here model Lamin A/C deficiency rather than normal aging, and aging involves many additional processes, including mitochondrial dysfunction, inflammation, stem-cell exhaustion, altered proteostasis, and systemic changes in tissue environment. Thus, our data should not be interpreted as showing that Lamin A/C disruption is the cause of aging. Rather, they support a more specific and justified idea: Lamin A/C-dependent mechano-epigenetic organization may represent one mechanism through which mechanically stressed tissues maintain nuclear and transcriptional homeostasis, and disruption of this mechanism may contribute to aging-like or degenerative features in certain contexts^28^.

Overall, this study advances the field by showing that the physiological function of Lamin A/C cannot be fully understood from nuclear shape in cultured cells alone. In intact tissues, Lamin A/C deficiency can leave the nuclear outline largely preserved while disrupting the mechanical and epigenetic organization of the nuclear interior. The live *in vivo* measurements show that Lamin A/C contributes to nuclear mechanical homeostasis in skeletal muscle, while the chromatin and transcriptomic data show that this mechanical role is coupled to heterochromatin organization and gene-expression buffering. We propose that Lamin A/C maintains a mechano-epigenetic state in which heterochromatin remains mechanically load-bearing, H3K9me3 remains spatially coupled to compact chromatin, and muscle transcriptional programs remain stable. Partial Lamin A/C disruption reveals a compensatory state in which chromatin architecture is altered but gene expression is buffered, whereas complete Lamin A/C loss causes failure of this compensation and broad myopathic transcriptional dysregulation. This framework connects Lamin A/C, nuclear mechanics, chromatin architecture, and transcriptional homeostasis *in vivo*, and provides a foundation for future studies of mechano-epigenetic regulation in muscle disease, tissue degeneration, and aging-related nuclear dysfunction.

## MATERIALS AND METHODS

### Lamin A/C-deficient mice

All animals experiments were performed after receiving the IACUC protocol at University of Colorado Boulder. B6129S1(Cg)-*Lmna^tm1Stw^*/BkknJ mice were purchased from Jackson Laboratory (#009125). This targeted allele expresses a truncated Lamin A/C protein that does not properly integrate into the nuclear lamina. Heterozygous mice (*Lmna*^+/-^) are viable and fertile and were bred to generate wild-type (*Lmna*^+/+^; WT), heterozygous (*Lmna*^+/-^; HZ), and homozygous Lamin A/C-deficient (*Lmna*^-/-^; LD) mice. LD mice exhibit severe postnatal growth retardation beginning around 2 weeks of age, abnormal movement and gait by 3–4 weeks of age, and typically die around 7-8 weeks of age. All mice were weaned at postnatal day 21 and housed in a temperature-controlled environment under a 12-h light/dark cycle, with food and water available ad libitum. Homozygous LD mice were additionally provided DietGel Recovery (Clear H2O) to support feeding and hydration. Mice were genotyped by tail biopsy according to the Jackson Lab protocol. For all experiments 6-7 week animals were used.

### Tissue collection for atomic force microscopy and immunofluorescence

Mice were euthanized by carbon dioxide overdose, and necropsy was performed to collect brain, skeletal muscle (Tibialis Anterior for microscopy and AFM, Quadriceps for ATAC seq and RNA seq), heart, liver, lung, cartilage (knee joint), and skin tissues. Fresh tissues were used for atomic force microscopy to quantify tissue stiffness. Except for cartilage, tissues were embedded in agarose and sectioned at 100–200 µm thickness using a vibratome. For optical microscopy, tissues were fixed overnight in 4% paraformaldehyde and then processed for DAPI, phalloidin, and immunofluorescence staining.

### Atomic force microscopy

Fresh tissue sections were used to quantify tissue stiffness by atomic force microscopy (Keysight 5500, Agilent). Elastic modulus was measured in skeletal muscle, heart, and brain tissue sections. Measurements were performed on tissue sections from four animals per group, and individual measurements were used to calculate group-level stiffness distributions.

### Neuromuscular stimulation and live in vivo measurement of multiscale mechanics

To quantify tissue-to-nucleus strain transfer in skeletal muscle *in vivo*, we used neuromuscular stimulation as described previously. Briefly, mice were anesthetized and placed in the prone position. The gastrocnemius muscle was surgically exposed, and nuclei were stained using NucBlue Live ReadyProbes Reagent (Thermo Fisher Scientific). The deep fibular nerve was stimulated to induce hindlimb muscle contraction, and skeletal muscle tissue (gastrocnemius) and nuclei were imaged before and during stimulation using an inverted confocal microscope (Nikon Eclipse Ti A1R) with 10× and long-working-distance 40× objectives. Confocal image pairs acquired before and during stimulation were used to quantify tissue-scale and nuclear-scale deformation. Mechanical strain inside the tissue and nucleus was calculated using deformation microscopy, which has been previously validated in our earlier works^30,31^. Briefly, undeformed and deformed images were registered using a hyperelastic warping algorithm to calculate displacement fields. These displacement fields were then used to compute hydrostatic, deviatoric, and shear strain maps in the tissue and nucleus.

### Immunofluorescence staining and imaging

Fixed tissue sections were permeabilized with 1% Triton X-100 in PBS for 1 h and blocked with 10% normal goat serum and 1% bovine serum albumin in 0.1% PBT, consisting of 0.1% Tween-20 in PBS, for 1 h. Primary antibody incubation was performed at 4 °C for 48 h in 0.1% PBT containing 1% bovine serum albumin. Primary antibodies were used against H3K9me3 (mouse monoclonal, Thermo Fisher MA5-42567), H3K27me3 (rabbit monoclonal, Thermo Fisher MA5-11198), H3K9ac (rabbit monoclonal, Thermo Fisher MA5-11195), Lamin A/C (mouse monoclonal, Cell Signaling Technology 4777), and Lamin B (rabbit polyclonal, Abcam ab16048). Secondary antibodies were goat anti-rabbit and anti-moue at different wavelengths (Thermo Fisher A32731, A32733 and A32727). Secondary antibody incubation was performed at room temperature for 3 h in the same buffer at a dilution of 1:200.

Some immunofluorescence images, including those used for super-resolution analysis, were acquired using a Zeiss LSM 980 microscope in Airyscan/super-resolution mode with a 63× oil-immersion objective. Other immunofluorescence images, including those used for tissue-level nuclear shape and chromatin architecture analysis, were acquired using a Nikon Eclipse Ti A1R inverted confocal microscope. Raw images were used for quantitative analysis. Skewness and kurtosis of DAPI or histone-mark intensity distributions were computed from regions of interest using Fiji. Nuclear shape parameters, including those used to calculate the nuclear irregularity index, were also measured in Fiji. Correlation analysis between multiple channels (DAPI and H3K9Me3) was performed using an in-house Matlab code.

### Nuclear shape and chromatin intensity analysis

Nuclear morphology was quantified from confocal images using Fiji. Nuclear regions of interest were segmented from DAPI images, and nuclear-shape parameters were measured for each nucleus. A nuclear irregularity index was calculated from multiple geometric parameters, including aspect ratio, radius ratio, circularity, and the ratio of nuclear area to bounding-box area. Higher values indicated greater deviation from a regular nuclear shape. Chromatin architecture was quantified using the spatial distribution of DAPI intensity within individual nuclei. DAPI-dense regions were interpreted as DNA-rich heterochromatin-like regions, whereas lower-intensity regions were interpreted as euchromatin-like regions. Skewness and kurtosis of intranuclear DAPI intensity distributions were calculated to quantify the degree of chromatin segregation. Higher skewness and kurtosis values indicated a more heterogeneous DNA-density landscape with stronger separation between dense and less dense chromatin regions. For H3K9me3 and other chromatin marks, intensity-distribution metrics were interpreted as measures of histone-mark spatial concentration and heterogeneity rather than direct measures of DNA compaction.

### Tissue collection and ATAC-seq

Quadriceps tissues were dissected from mice and processed in a single standardized tissue-preparation batch to minimize technical variability across samples. Immediately after isolation, tissues were flash-frozen in liquid nitrogen and stored at −80 °C until analysis. Frozen quadriceps samples were shipped on dry ice to Active Motif (Carlsbad, CA) for ATAC-seq. At Active Motif, nuclei were isolated from frozen tissue and subjected to Tn5 transposase-mediated fragmentation to tag accessible chromatin regions. Barcoded sequencing libraries were prepared using Active Motif’s validated ATAC-seq workflow, followed by library quality control, quantification, and Illumina paired-end sequencing. Active Motif performed initial bioinformatic processing, including read trimming, alignment to the mouse genome (mm10), peak calling, and generation of quality-control metrics. FASTQ files, aligned BAM files, peak calls, bigWig coverage tracks, and differential accessibility outputs were used for downstream chromatin-accessibility analyses. Because ATAC-seq was performed with one biological sample per genotype, these data were interpreted as exploratory and hypothesis-generating. ATAC-seq results were used to support and extend imaging-based observations of chromatin remodeling rather than as the primary basis for the central conclusions.

### Bulk RNA-sequencing

Quadriceps tissues were dissected, rinsed in DPBS, flash-frozen, and stored at −80 °C until processing by Novogene. RNA-seq was performed with n = 4 animals per group. RNA-seq analysis was performed using Partek software (Partek Inc., St. Louis, MO). Raw sequence reads were first subjected to quality-control checks and then aligned to the mouse genome (mm10) using STAR. Quality checks were performed on aligned reads, and HTSeq was used to quantify gene-level counts. Genes with zero counts in at least 80% of samples were filtered out. Count data were normalized using the median-ratio method, and DESeq2 was used for differential expression analysis between sample groups.

### Integrated ATAC-seq and RNA-seq analysis

Promoter-associated differentially accessible regions were compared with differentially expressed genes to evaluate the relationship between chromatin accessibility and gene expression across genotypes. ATAC-seq peak annotations were used to associate accessible regions with nearby genes, and RNA-seq differential expression results were used to compare chromatin-accessibility changes with transcript-level changes. Candidate loci, including *Hdac2* and *Cdkn1a*, were further examined using ATAC-seq track visualization and RNA-seq expression data.

### Gene ontology and motif enrichment analysis

Gene Ontology biological process enrichment analysis was performed on differentially expressed genes and genes associated with differentially accessible regions. Motif enrichment analysis was performed on differentially accessible chromatin regions to identify transcription factor binding motifs associated with genotype-dependent accessibility changes.

### Statistical analysis

The number of biological and technical replicates is reported in each figure or figure caption. Plots were prepared using Microsoft Office, R, or other indicated software. Bar graphs show mean ± standard deviation unless otherwise stated. Where applicable, one-way ANOVA was performed followed by Tukey’s post hoc test. Specific statistical tests are reported in figure captions.

## Acknowledgements

This work was supported by NSF CAREER 1349735 and 2212121 to C.P.N., and by NSF CAREER 2236710 and NSF 2322878 to S.G.

## Author contributions

S.G. and C.P.N. conceptualized the study; S.G., K.H., A.K.S, S.E.S., B.S. and C.P.N planned and implemented the methodology; software, S.G., K.H.; formal analysis, S.G., K.H., J.K., N.C.; investigation, S.G., S.E.S., A.K.S., K.H., X.X., B.S.M.M.; S.G. wrote the original draft; all authors reviewed and edited; S.G. and C.P.N. acquired funding; S.G. and C.P.N. provided the resources.

## SUPPLEMENTARY FIGURES

**Figure S1.**
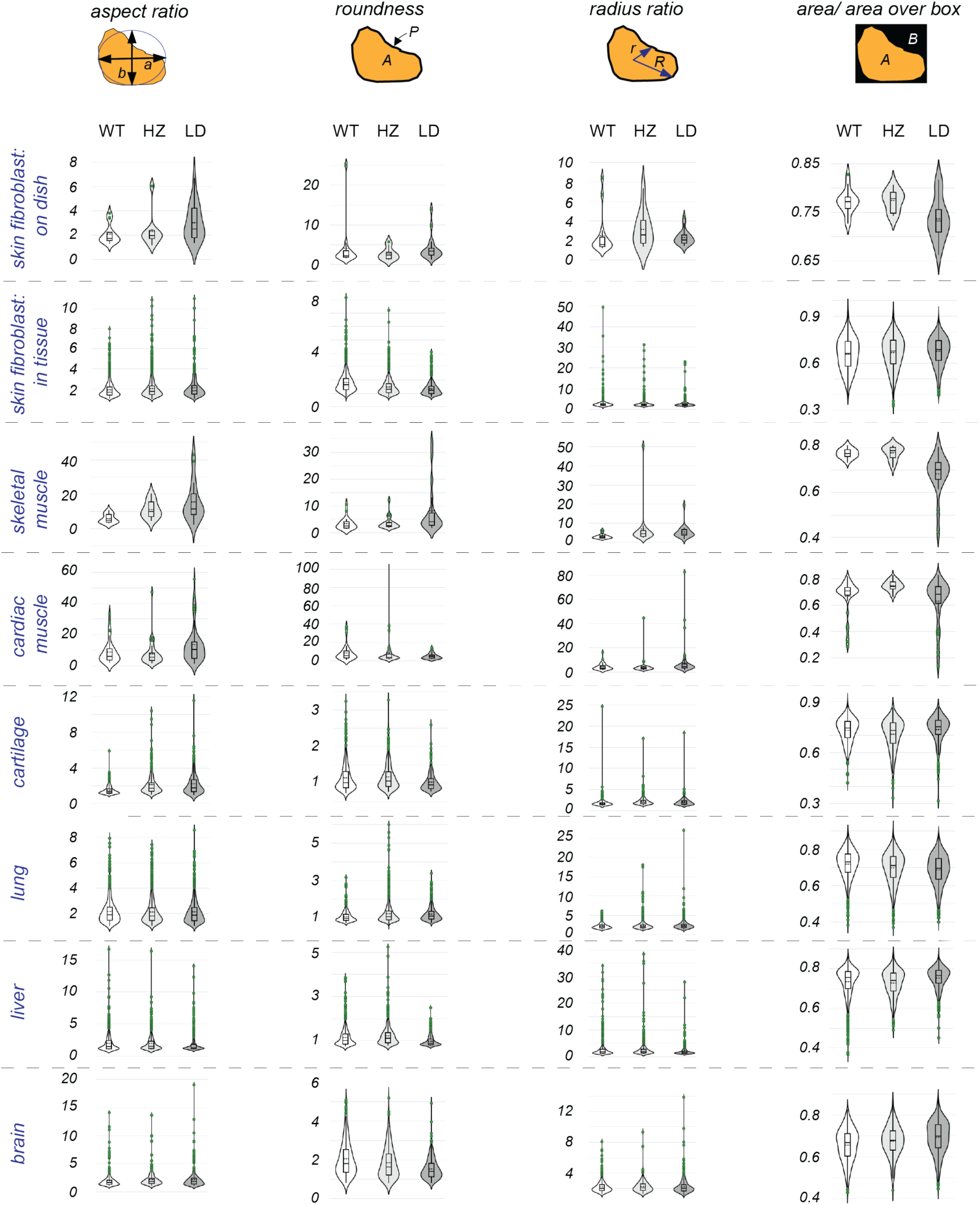
Nuclear irregularity index used to quantify nuclear shape. The nuclear irregularity index used in Figure 1 was calculated from multiple geometric parameters describing nuclear shape: aspect ratio, radius ratio, circularity, and the ratio of nuclear area to bounding-box area. The index was defined as: nuclear irregularity index = aspect ratio + radius ratio + circularity - 1 - nuclear area/bounding-box area. Higher values indicate greater deviation from a regular nuclear shape. Violin plots show the full distribution of nuclear irregularity values, with embedded boxplots and dotted lines indicating the mean.

**Figure S2.**
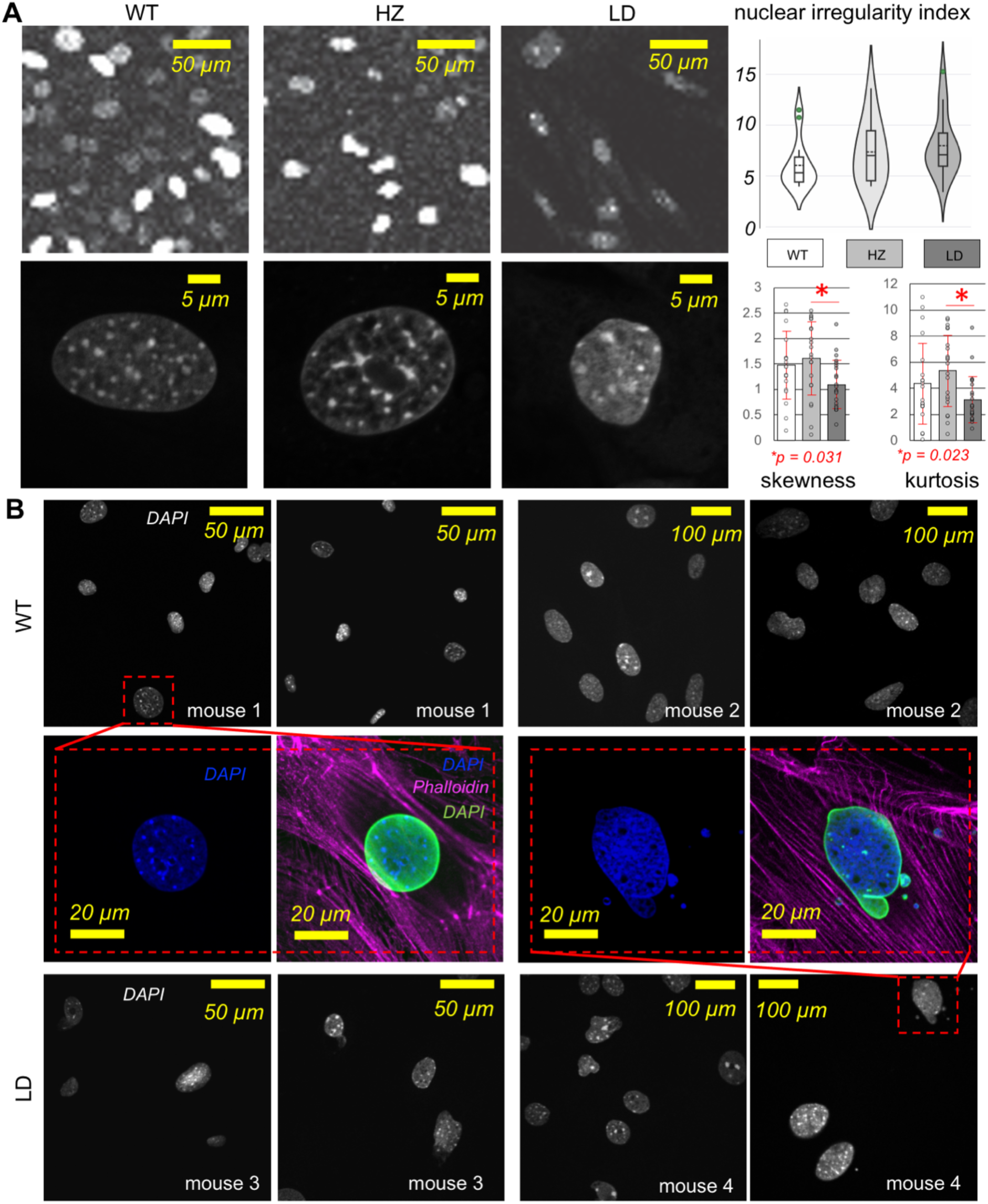
Lamin A/C-deficient fibroblasts show nuclear-shape abnormalities in 2D monolayer culture in vitro. **(A)** Representative images of skin-derived fibroblasts cultured as 2D monolayers. A subpopulation of LD fibroblast nuclei showed irregular shape compared to WT and HZ nuclei, which were more commonly round and regular. Higher-resolution images show nuclear-shape abnormalities in LD fibroblasts. Although the phenotype was visually apparent in some cells, the difference was not statistically significant in this dataset, based on one mouse per group and >50 nuclei per mouse. DAPI intensity analysis also suggested more diffuse chromatin organization in LD fibroblasts, indicated by lower skewness and kurtosis. **(B)** Cardiac fibroblasts showed a similar trend. Even in WT cultures, some nuclei were irregular, but LD cultures contained a larger subpopulation of nuclei with abnormal shape and occasional blebbing. These data confirm that 2D monolayer culture can reveal the expected Lamin A/C-dependent nuclear-shape phenotype, while also showing that the phenotype is heterogeneous across cells.

**Figure S3.**
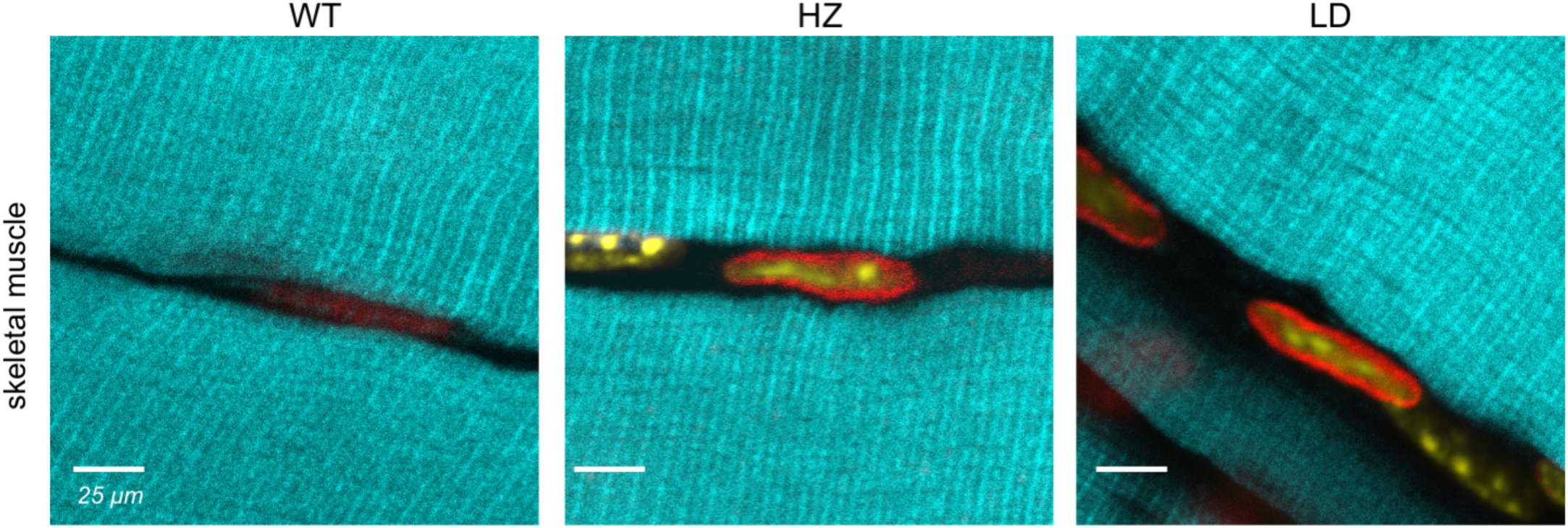
Lamin A/C deficiency does not visibly disrupt skeletal muscle tissue architecture. Supplementary images corresponding to Figure 2 showing skeletal muscle tissue architecture in WT, HZ, and LD mice. Phalloidin staining marks F-actin (cyan), DAPI marks nuclei (yellow), and Lamin B marks the nuclear envelope (red). Skeletal muscle fibers retained gross tissue organization across genotypes, supporting the conclusion that Lamin A/C deficiency does not produce a broad collapse of tissue architecture in intact skeletal muscle.

**Figure S4.**
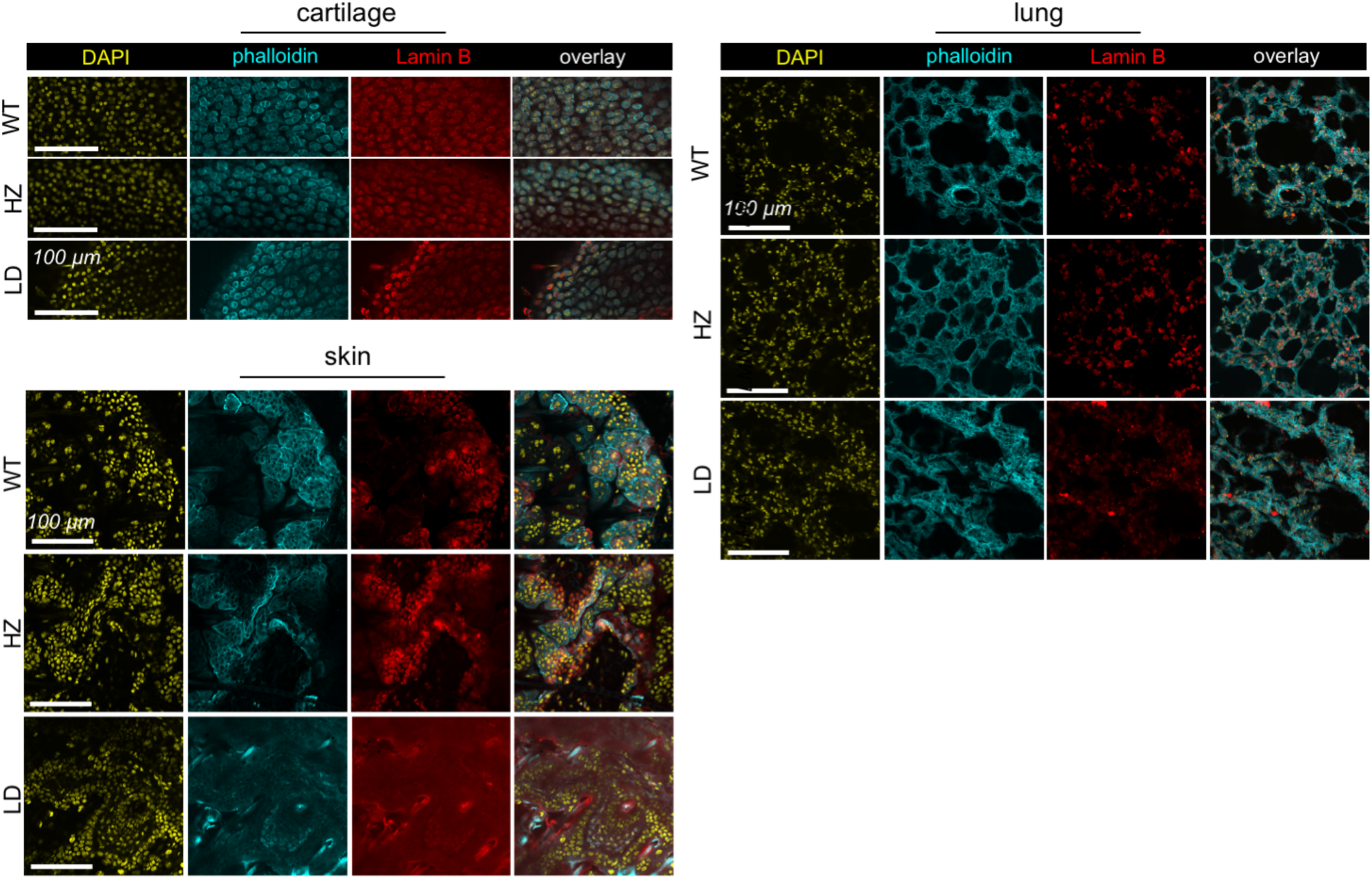
Lamin A/C deficiency does not visibly disrupt tissue architecture in cartilage, lung, and skin. Supplementary images corresponding to Figure 2 showing cartilage, lung, and skin tissues from WT, HZ, and LD mice. Phalloidin and nuclear staining revealed no obvious genotype-dependent disruption of gross tissue architecture in these tissues. These data support the conclusion that preserved tissue organization may contribute to the maintenance of nuclear shape in vivo.

**Figure S5.**
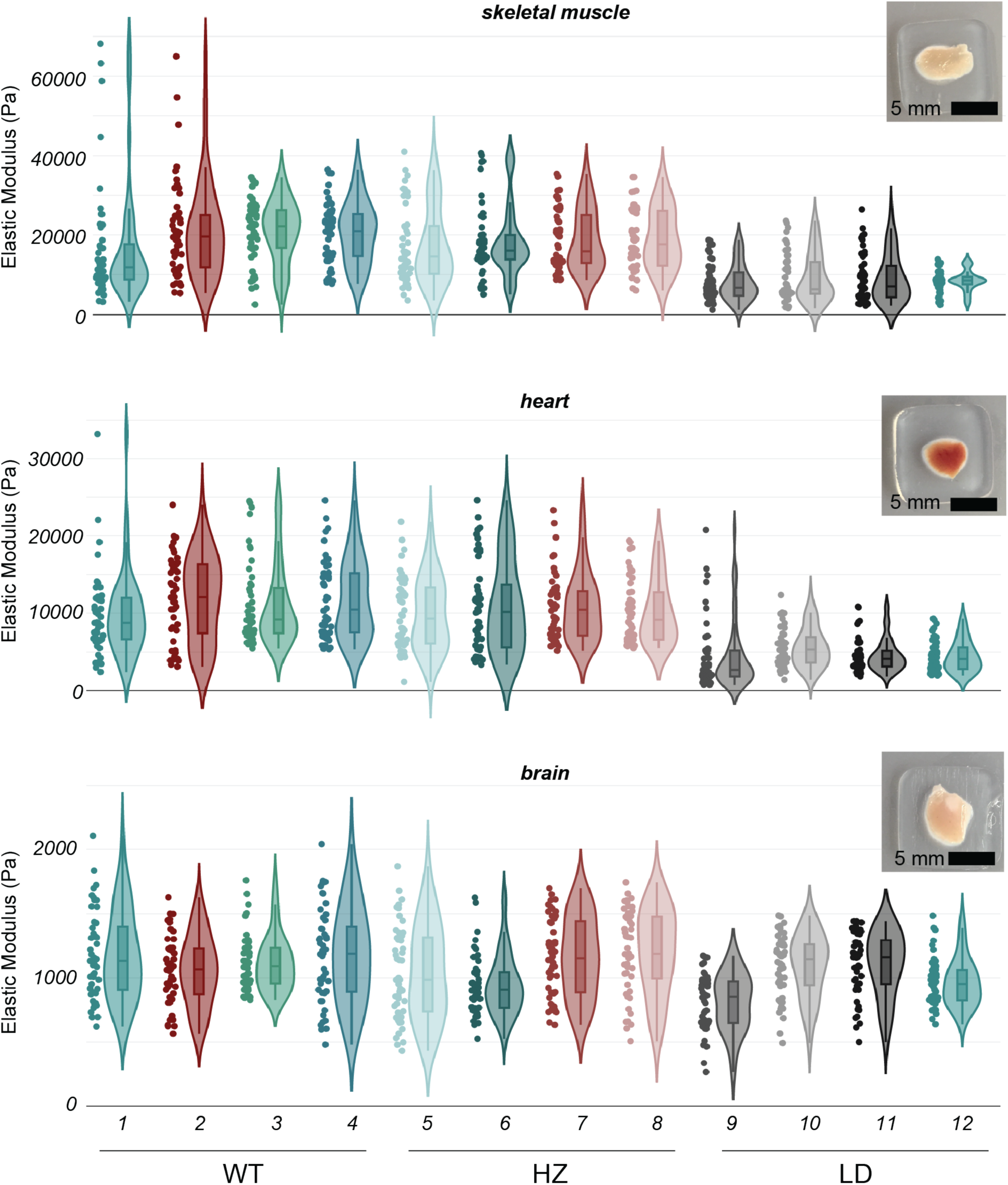
Atomic force microscopy quantification of tissue stiffness. Atomic force microscopy measurements of elastic modulus in tissue sections from skeletal muscle (tibialis anterior), heart, and brain (see Figure 4D). Data show individual measurements from four animals per group. Lamin A/C deficiency reduced tissue stiffness in skeletal muscle and heart, whereas brain stiffness was comparatively unchanged.

**Figure S6.**
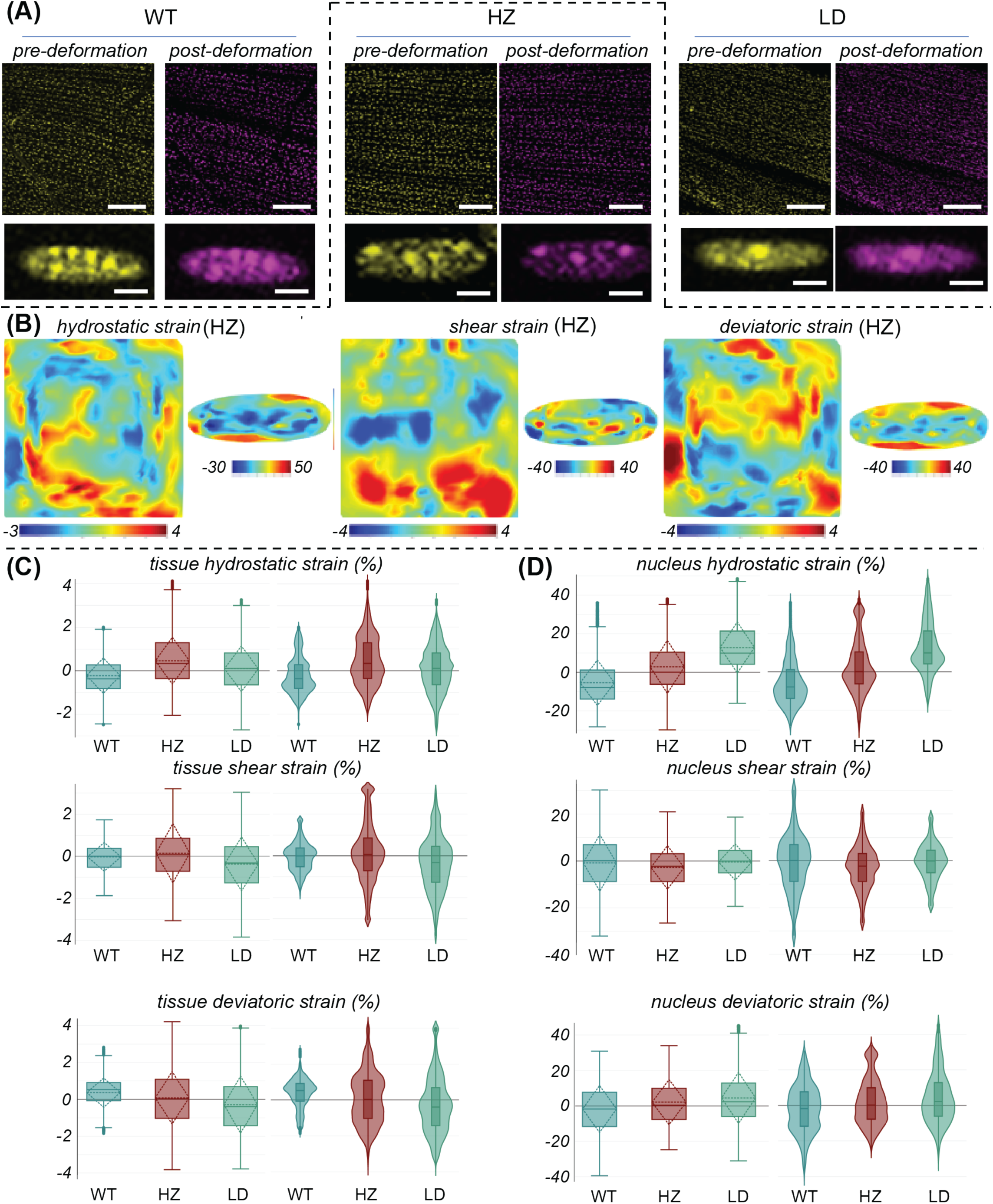
In vivo multiscale deformation analysis of skeletal muscle tissue and nuclei. Supplementary data corresponding to Figure 4. **(A)** Representative undeformed and deformed confocal images of skeletal muscle tissue and nuclei from WT, HZ, and LD mice acquired during live in vivo deformation experiments. **(B)** Hydrostatic, deviatoric, and shear strain maps for the LD group, showing spatially heterogeneous deformation at both tissue and nuclear scales. **(C)** Distribution of tissue-level hydrostatic strain for the representative sample shown in panel A. **(D)** Distribution of nuclear hydrostatic strain for the same representative sample. These data illustrate how deformation microscopy was used to quantify multiscale strain transfer from tissue to nucleus in living skeletal muscle.

**Figure S7.**
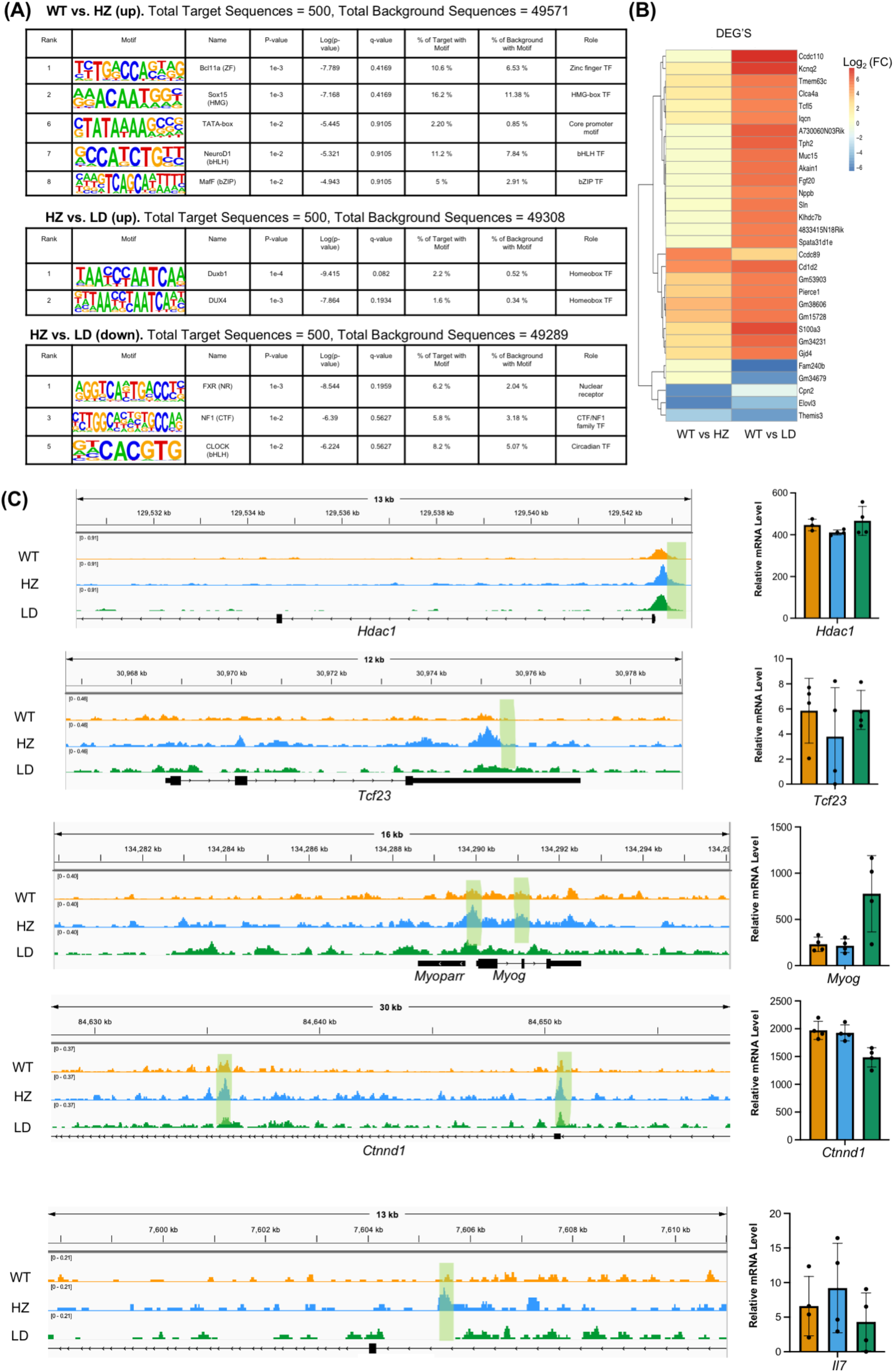
Additional ATAC-seq and RNA-seq analyses of Lamin A/C-deficient skeletal muscle. **(A)** Motif enrichment analysis of differentially accessible regions in WT versus HZ, HZ versus LD, and additional pairwise comparisons. These analyses provide further insight into transcription factor motifs associated with chromatin-accessibility changes after partial or complete Lamin A/C disruption. **(B)** Heatmap showing highly differentially expressed genes across WT, HZ, and LD groups, highlighting genotype-dependent transcriptomic changes. **(C)** ATAC-seq tracks and relative mRNA levels for Hdac1 and Tcf23. WT: orange, HZ: blue, LD: green. Hdac1 showed accessible chromatin regions across groups with relatively consistent mRNA expression. Tcf23, a negative regulator of myogenic differentiation, showed greater accessibility in HZ but not WT or LD, with modestly lower relative mRNA levels in HZ. These data further illustrate that chromatin accessibility and gene expression are not globally coupled in a simple linear manner.

**Figure S8.**
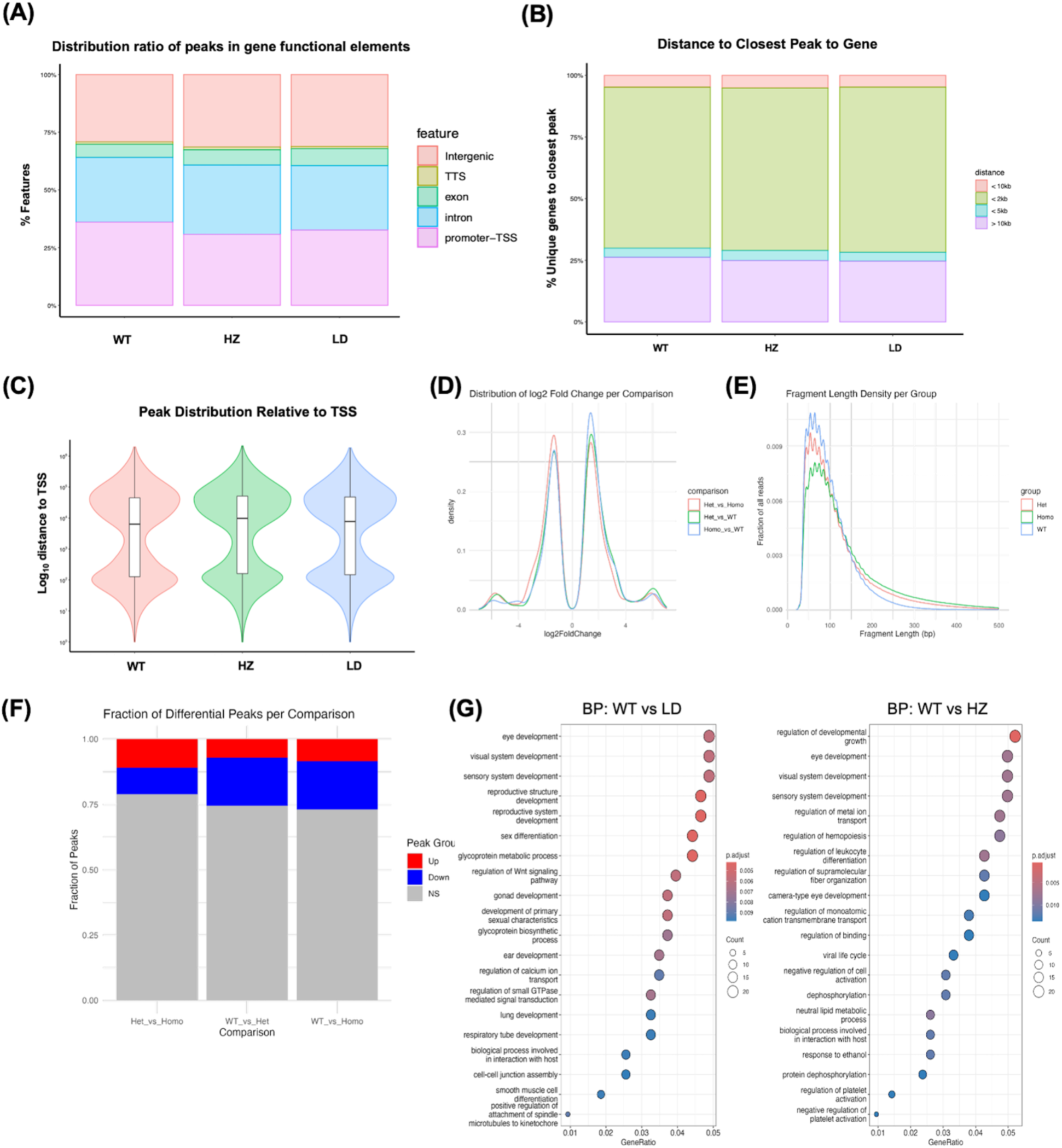
Quality-control metrics for ATAC-seq analysis. Quality-control analyses supporting ATAC-seq profiling of WT, HZ, and LD quadriceps muscle. **(A–B)** Distribution of peaks across gene functional elements and distance of peaks to transcription start sites. HZ showed the lowest percentage of promoter-associated peaks, whereas WT showed the highest, while peak-distance distributions were broadly comparable across groups. **(C–E)** Additional quality-control metrics generated by Active Motif, including peak distribution relative to transcription start sites, peak density, and fragment-length distribution. These metrics showed broadly consistent data quality across samples. **(F)** Differential accessibility analysis showing the fraction of peaks that were increased, decreased, or not significantly changed across pairwise comparisons. HZ versus LD contained the largest fraction of non-significant peaks, suggesting partial similarity between these chromatin-accessibility states. **(G)** Additional Gene Ontology pathway analysis for WT versus LD and WT versus HZ comparisons, supporting the involvement of muscle-related, signaling, and regulatory pathways in Lamin A/C-dependent chromatin remodeling.

**Figure S9.**
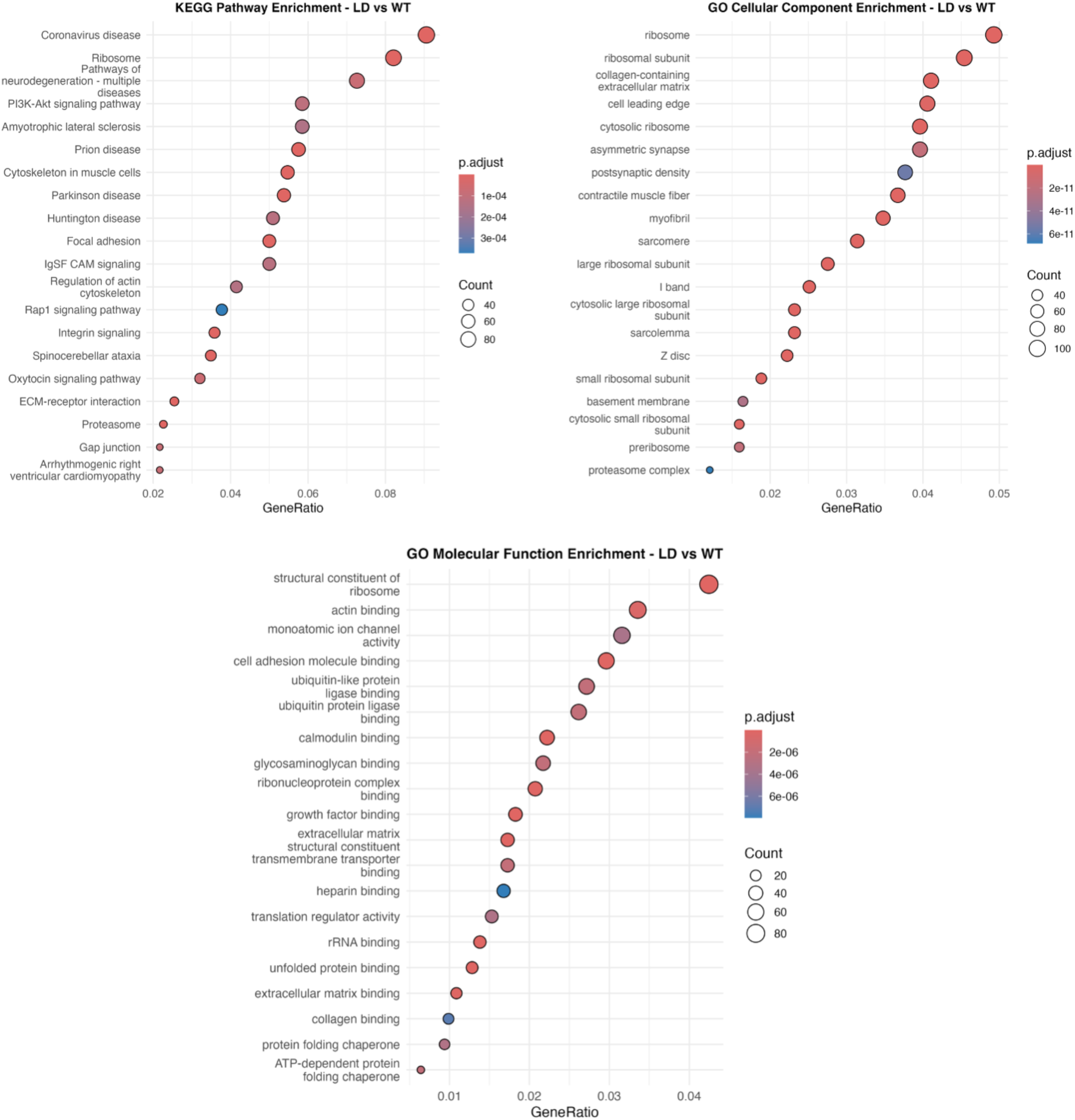
Pathway enrichment analysis between WT and LD groups based on RNA seq data in muscle. Specific molecular biological and functional pathways are more prominently or less prominently enriched in Lamin deficient mouse compared to wild type mice. Data based on RNA sequencing of quadriceps muscle (n = 4 animals/ group).

